# Sex-specific phenotypic effects and evolutionary history of an ancient polymorphic deletion of the human growth hormone receptor

**DOI:** 10.1101/788653

**Authors:** M. Saitou, S. Resendez, A.J. Pradhan, F. Wu, N.C. Lie, N.J. Hall, Q. Zhu, L. Reinholdt, Y. Satta, S. Nakagome, N.A. Hanchard, G. Churchill, C. Lee, G.E. Atilla-Gokcumen, X. Mu, O. Gokcumen

## Abstract

The deletion of the third exon of the growth hormone receptor (*GHR*) is one of the most common genomic structural variants in the human genome. This deletion (*GHRd3)* has been linked to response to growth hormone, placenta size, birth weight, growth after birth, time of puberty, adult height, and longevity. However, its evolutionary history and the molecular mechanisms through which it affects phenotypes remain unresolved. Here, we analyzed thousands of genomes and provide evidence that this deletion was nearly fixed in the ancestral population of anatomically modern humans and Neanderthals. However, it underwent a paradoxical adaptive reduction in frequency approximately 30 thousand years ago in East Asia that roughly corresponds with the emergence of archaeological evidence for multiple modern human behaviors, dramatic changes in climate, and a concurrent population expansion. Further, we provide evidence that GHRd3 is associated with protection from edematous severe acute malnutrition primarily in males. Using a mouse line engineered to contain the deletion, we found *Ghrd3*’s effect on the liver transcriptome of male mice grown without any calorie restriction mimics response to calorie restriction through regulation of circadian pathways. In contrast, under calorie restriction, *Ghrd3* leads to the female-like gene expression in male livers. As a likely consequence, the dramatic weight difference between male and female mice disappears among *GHRd3* mice under calorie restriction. Our data provide evidence for sex- and environment-dependent effects of *GHRd3* and are consistent with a model in which the allele frequency of *GHRd3* varies throughout human evolution as a response to fluctuations in resource availability.

---

Why does functional genetic variation remain polymorphic? This question is fundamental to evolutionary biology. The complete deletion of the third coding exon of the growth hormone receptor gene (*GHR*) in humans provides a compelling model to study this question. This 2.7kb deletion (*GHRd3;* esv3604875) is found between 10% to 80% allele frequency among human populations and in all sequenced Neanderthal and Denisovan genomes. This observation is unexpected given that the receptor is highly conserved among mammals and fundamental in various cellular processes, including cell division, immunity, and metabolism (*1*), and a loss of function mutation of this gene causes recessive Laron syndrome (*2, 3*). The *GHRd3* allele generates a shorter isoform of this critical developmental gene (*1*). Locus-specific studies consistently and reproducibly associate *GHRd3* with altered placental and birth weight (*4*), time of puberty onset (*5*), lifespan (*6*), metabolic activity (*7*), and response to growth hormone treatments (*8*). Despite the known relevance of *GHRd3* to human phenotypes that are likely crucial for human evolution, the mechanisms through which *GHRd3* affects cellular and organismal function and the evolutionary forces that maintain the *GHRd3* allele in human populations remain mostly unknown.

*GHRd3* has a complex evolutionary history in humans. Our previous work determined that *GHRd3* is one of 17 exonic, polymorphic human deletions shared with the Altai Neanderthal or Denisovan genomes (*9*). This study now extends this observation to include the Vindija (*10*) and Chagyrskaya (*11*) Neanderthal genomes (**Fig. S1A**). The fact that this derived deletion is shared with archaic humans suggests that the *GHRd3* allele was formed before the divergence of the human and Neanderthal lineages. Further, while the deletion remains polymorphic in humans, it was likely fixed in the Neanderthal and Denisovan lineages since all of the 4 archaic human genomes we analyzed carry the deletion allele homozygously. We dismissed Neanderthal introgression as a possible explanation for the allele sharing since African populations have the highest allele frequency for this deletion (*9*). The deleted coding sequence is conserved among mammals (**Fig. S1B**). On a broader scale, exon 3 is significantly more conserved than randomly selected sites on the same chromosome (p = 1.3x10^-8^, Kolmogorov–Smirnov test), suggesting that this exon is not evolving under neutral conditions (**Fig. S1C**). When these data are observed in conjunction with the existing literature associating *GHRd3* with multiple human traits, it raises the possibility that adaptive forces may have shaped the allele distribution of this variant among extant human populations.

To understand the evolutionary forces that have maintained the *GHRd3* allele in human populations, we resolved the linkage disequilibrium (LD) architecture around *GHRd3* (**Fig. S2**). The deletion is in LD (r^2^ > 0.75) with fifteen nearby single nucleotide polymorphisms (SNPs) (**Table S1**). These variants help construct a short haplotype block (Hg19, chr5: 42625443-42628325) flanked by known recombination spots. This specific haplotype tags the *GHRd3* allele. Based on these tag-SNPs and direct genotyping of the deletion available through the 1,000 Genomes dataset, we found that *GHRd3* is the major allele in most African populations (**Fig. 1A, Fig. S3**). However, the allele frequency decreases to less than 25% in most Eurasian populations. The single nucleotide variants with strong LD with the deletion allowed for the construction of a haplotype network where one can visualize the clear separation of haplotypes that carry the *GHRd3* allele from those that do not (**Fig. 1B**). One unexpected observation is that the haplotypes that harbor the deletion are more variable and cluster with the chimpanzee haplotype than those that do not (**Fig. S4**). In other words, the ancestral non-deleted allele is harbored primarily by recently evolved haplotypes. This observation raises the possibility that *GHRd3* was nearly fixed in the ancestral human lineage and that the haplotypes that harbor the ancestral, non-deleted allele rebounded in frequency only recently.

**Figure 1.**
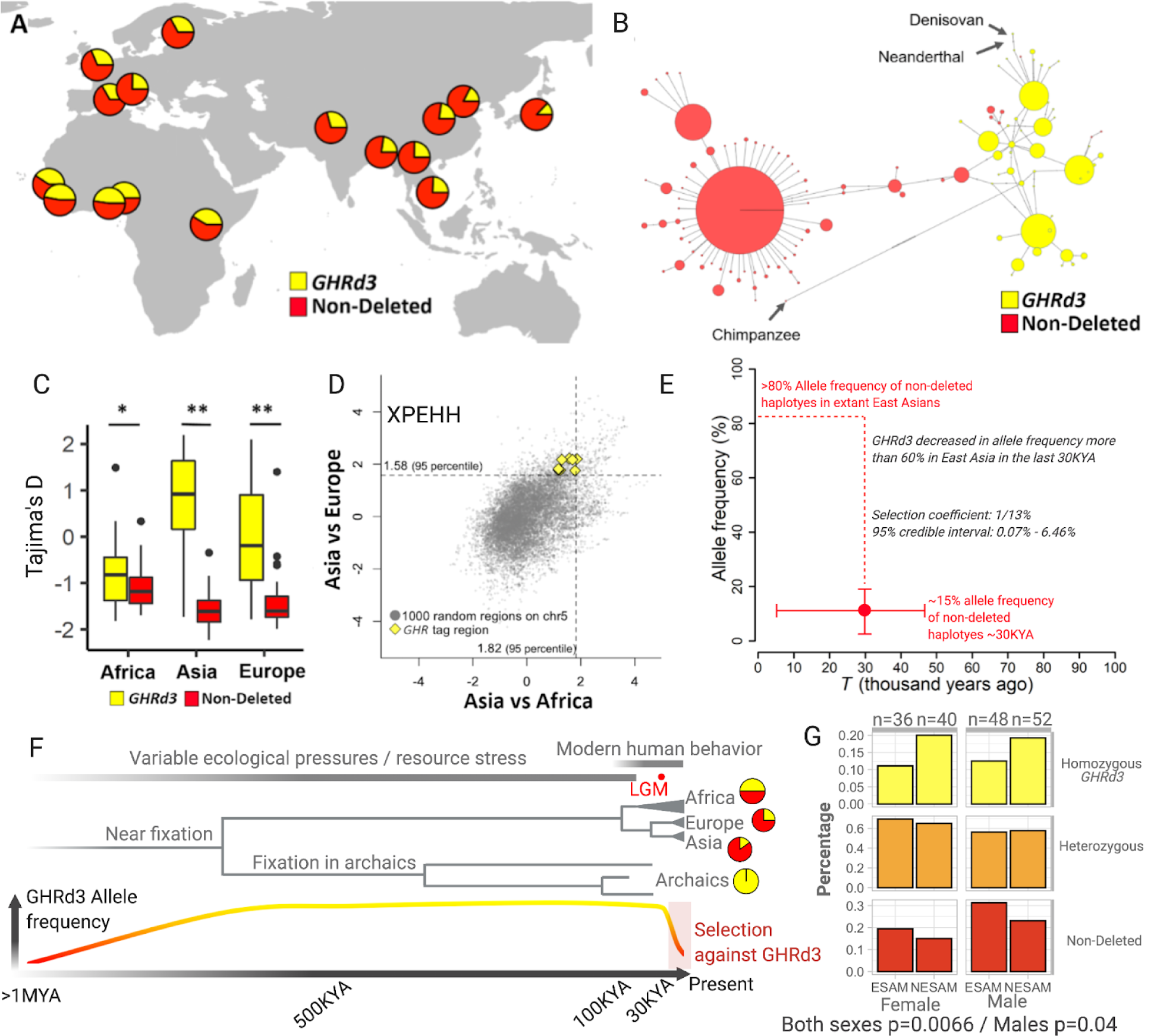
**A.** The geographic distribution of the *GHRd3* polymorphism. The deletion status is color-coded for the entire figure. **B.** A network of 2,504 human haplotypes, the Altai Neanderthal genome, the Denisovan genome, and the chimpanzee reference genome, calculated from variations within the high LD region upstream of the *GHRd3* location (Hg19: chr5: 42624748-42628325, see **Fig. S1**). **C.** Tajima’s D values are calculated for the deleted and non-deletion alleles in three 1000 Genomes meta-populations. The * and ** indicate 0.05 and 0.01 significance, respectively. **D.** XP-EHH values calculated for the *GHRd3* upstream region.; XP-EHH values are computed for the *GHRd3* tag SNPs (r^2^ > 0.8) and compared to distributions calculated for 1000 similarly sized regions (shown in grey). **E.** Estimation of the allele frequency of non-deleted haplotypes (and by proxy that of *GHRd3*) and the date for the onset of selection in the CHB population based on the results of the ABC simulations. The x-axis shows time, while the y-axis represents allele frequency. This plot shows means of *T* and *f_t_* with 95% credible intervals. The dotted line indicates the difference in allele frequency between the present and *t*. **F.** A plausible model for the evolution of *GHRd3.* From bottom to top, we indicated GHRd3 allele frequency across time where yellow depicts deleted allele and red depicts non-deleted allele. Next, we show the phylogenetic relationships between extant and archaic humans with the allele frequencies shown with the pie charts at the branch tips. The timeline for ecological variability and the onset of modern human behavior was shown at the top of this panel. LGM: Last Glacial Maximum. **G.** The association between *GHRd3* and edematous severe acute malnutrition (SAM). GHRd3 is significantly enriched in non-edematous SAM as compared to edematous SAM, indicating that GHRd3 may provide protection against developing the classically more severe form of acute malnutrition. The graph shows the percentages of Non-Deleted, heterozygous, and homozygous GHRd3 genotypes in each category. The sample size of each category is indicated on the top of the chart.

**Table 1.** The effect sizes of genes that show significant expression level differences between wt/wt and d3/d3 mice in AL and CR conditions. The orange and blue colors indicate genes that are upregulated or downregulated more than 2 fold in d3/d3 mice compared to wt/wt mice, respectively. The fold-changes that are shown to be significant (p_adj_ < 0.01) are bolded.

| Gene | Effect of Ghrd3 under AL | Effect of Ghrd3 under CR | Significant in |
| --- | --- | --- | --- |
| Steap4 | **-1.905329326** | -0.41 | AL |
| Tef | **0.958439066** | 0.14 | AL |
| Pias3 | **0.572051345** | 0.06 | AL |
| Per3 | **1.344894349** | 0.22 | AL |
| Etv6 | **-0.540772517** | -0.06 | AL |
| Mt1 | **-2.939960534** | 0.32 | AL |
| Tle3 | **-0.984835227** | 0.59 | AL |
| Ppargc1b | **1.173267184** | -0.23 | AL |
| Egr1 | **-3.728386881** | -0.68 | AL |
| Ciart | **2.489709512** | -0.39 | AL |
| Dbp | **2.793636017** | -0.28 | AL |
| Alyref2 | **-0.741847732** | -0.02 | AL |
| Cry2 | **0.679940266** | 0.13 | AL |
| Tspan33 | -0.168515679 | **-0.76** | CR |
| C9 | -0.114731926 | **-2.21** | CR |
| C6 | -0.036758302 | **-1.62** | CR |
| Arsa | 0.234501888 | **-0.85** | CR |
| Mcm10 | -0.422994919 | **-1.36** | CR |
| Fmo3 | -0.170930379 | **2.64** | CR |
| Styk1 | -0.248481672 | **-1.86** | CR |
| Hsd3b5 | -0.720935552 | **-3.86** | CR |
| Susd4 | 0.179893878 | **-1.54** | CR |
| Rsph4a | 1.072477146 | **1.89** | CR |
| Cyp2b13 | -0.779783638 | **4.44** | CR |
| Slco1a1 | -0.094227348 | **-2.44** | CR |
| Insc | -0.50623682 | **-1.03** | CR |
| Slc22a26 | **-2.093943342** | **2.08** | CR |
| Serpina3k | -0.190141472 | **-0.99** | CR |
| Serpina9 | **-1.915370404** | **-4.45** | CR |
| Ugt2b38 | -0.418122161 | **-2.76** | CR |
| Ces3b | -0.148815361 | **-1.20** | CR |
| Mup21 | -0.416888474 | **-2.72** | CR |
| Sult3a1 | 1.953997451 | **4.00** | CR |
| Ces3a | -0.117609056 | **-1.13** | CR |
| Mup7 | -0.088474963 | **-2.54** | CR |
| Cyp2a4 | 0.315997716 | **2.26** | CR |
| Gm13597 | -0.883393905 | **6.50** | CR |
| Mup20 | -0.02271503 | **-1.72** | CR |
| Serpina4-ps1 | **-2.42000237** | **-5.00** | CR |
| Rnf170-ps | 0 | **5.34** | CR |
| Gm29920 | 0 | **6.30** | CR |

To test if recent selection is acting on this locus, we used the single nucleotide variation data within the *GHRd3* upstream haplotype block to calculate Tajima’s D (*12*) and XP-EHH (*13*) values. These statistics measure deviations from expected allele frequency spectra (Tajima’s D) and extended homozygosity differences between populations (XP-EHH) under neutrality. We observed a significantly lower Tajima’s D value for the *GHR* locus in the East Asian population when compared to randomly selected regions on chromosome 5 (p < 0.05, Mann-Whitney (MW) test, **Fig. S5**). Moreover, the non-deleted haplotypes showed lower Tajima’s D values when compared to the deleted haplotypes in the Eurasian populations (p < 0.01, MW test, **Fig. 1C**). Concordant with the Tajima’s D analysis, when the *GHRd3* locus of the Han Chinese (CHB) population was compared with that of the West European (CEU) and Yoruba (YRI) populations, we noted XP-EHH values that were higher than genome-wide expectations (**Fig. 1D**). Collectively, these results are parsimonious with a relatively recent sweep of the existing non-deleted haplotypes in the East Asian population.

To summarize these results, a plausible model is that *GHRd3* was nearly fixed in the human-Neanderthal lineage and remained the major allele throughout most of human history, but only recently reduced in allele frequency due to relaxation of the selection favoring *GHRd3.* This scenario would present itself as a sweep on the existing haplotypes harboring the non-deleted allele. To investigate this model, we used the *Fc* method that combines site frequency spectrum analyses with LD calculations (*14*) and found a significant deviation of the Fc value from the simulated neutral expectations in the East Asian population, showing selection starting approximately 27 thousand years ago (Kolmogorov-Smirnov test, p < 0.01, **Supplementary Material**). To strengthen this analysis, we employed an approximate Bayesian computation method that compares the fit of different evolutionary models to the site frequency spectrum and haplotype structure variation data of a given locus (*15*) (**Fig. S6, Supplementary Material**). This approach selected a model that fits our hypothesis and suggested a recent sweep of the non-deleted haplotypes in East Asia. Specifically, it dated the onset of this sweep to 29,811 years ago, while putting the frequency of the non-deleted allele at ∼11% before the sweep (log_10_-scaled approximated Bayes factor, log_10_(aBF) = 5.250 against neutral; log_10_(aBF) = 13.257 against hard-sweep) (**Fig. 1E**). Collectively, the evolutionary analyses of this locus render neutral evolution a less likely explanation for the high allele frequencies of *GHRd3* and instead point to population-specific adaptive forces that likely vary across time.

As described earlier, *GHRd3* has been associated with multiple human traits. However, these associations were found in locus and phenotype-specific studies, often prone to type I errors. Moreover, most genome-wide association studies (GWAS), which focus primarily on single nucleotide variants, have not detected any associations between *GHRd3* and phenotypic variation. We argue that the inability of GWAS to pick up signals from this deletion is two-fold. First, SNP-array platforms, which rely on imputation to genotype *GHRd3*, interrogate a very limited number of variants in the locus. For example, Affymetrix 6.0 and Illumina OMNI Quad harbor only 4 and 3 SNPs flanking the deletion, respectively, with only 1 of those SNPs in both platforms having >95% LD with *GHRd3* (**Table S1**). Second, previous studies have shown that the effect of *GHRd3* is likely environment and sex-specific. As a result, the signal for *the GHRd3* association may be diluted in traditional GWAS.

To specifically search for the associations of *GHRd3* on human phenotypes, we examined the GWAS database (available through https://atlas.ctglab.nl/PheWAS) for phenotype associations using a single nucleotide variant that tags the deletion (rs4073476). Using a strict nominal *p*-value threshold of p < 10^-8^ in a phenome-wide association study (PheWas) of 152 phenotypes, we found that the derived *GHRd3* haplotype is strongly associated with bone mineral density. The next most significant association was height with a nominal p-value < 10^-5^ (**Table S1**). We then conducted a similar analysis in the UK Biobank dataset phenotypes (available through http://geneatlas.roslin.ed.ac.uk) (*16*). We found that *the GHRd3* haplotype is associated with standing height in this cohort (nominal p < 10^-8^, in a PheWAS of 742 phenotypes, **Table S1, Fig. S7**). These findings are concordant with previous locus-specific studies that link *GHRd3* with developmental and metabolic traits (*17*). Notably, one such locus-specific study underlined the sex-specificity of the phenotypic effect of *GHRd3* (*6*). Indeed, we found that among the top 10 traits that are at least nominally associated with *GHRd3* in the UK Biobank dataset, grip strength has a greater correlation with the deletion in males (p-values - left hand: 1.29 x 10^-5^; right hand: 1.40 x 10^-4^) than in females (p-values - left hand: 3.57 x 10^-1^; right hand: 1.48 x 10^-1^). This effect was noted independently for both the left and right hands, increasing our confidence in this observed trend.

Based on our evolutionary genetics insights and the predicted functional effect of *GHRd3* on metabolic and developmental traits, we hypothesize that this deletion may affect fitness within the context of the famine and abundance periods that have been a defining feature of human evolution (**Figure 1F**). It was previously speculated that the *GHRd3-*mediated upregulation of the growth hormone pathway might be a response to nutritional deprivation (*18*). Response to starvation is relevant to human evolution because compared to nonhuman great apes, humans generally cope with higher levels of seasonality and unpredictability of resource levels (*19*). In this ecological context, the fluctuation of resources may have been a major adaptive stressor for humans, especially on traits pertaining to metabolism and reproduction. Indeed, studies in other animals suggest that small size at birth and early reproduction, both noted effects of *GHRd3*, are favored under environmental stress (*20*). To further test this notion, we genotyped *GHRd3* in 176 Malawian children with severe acute malnutrition (SAM) (*21*) (**Supplementary Material**). We found that *GHRd3* is depleted among children who suffer from the more severe, edematous form of (SAM) (kwashiorkor) (Chi-Square test, p=0.0066), particularly in males (p=0.04, Chi-Square test) (**Figure 1G**). Thus, individuals carrying *GHRd3* may fare better under nutritional stress, supporting the hypothesis that *GHRd3* may confer a fitness advantage under such conditions. It is noteworthy that previous work showed that children who suffer from edematous SAM have significantly higher birth weight than those who suffer from less severe non-edematous (SAM) (*22*). Thus, it is plausible that lower birth weight at birth, which *GHRd3* is significantly associated with (*4*), may lead to a more favorable metabolic response to malnutrition and may increase fitness under nutritional stress.

Putting all this together, it is of note that *GHRd3* emerged 1-2 Million years ago. This emergence time coincides with the spread of the *Homo genus* within and outside of Africa (*23*), likely carrying the deletion across the world. It is plausible that *GHRd3* has increased the fitness of these archaic populations who were facing new ecologies and potential resource fluctuations. This scenario also fits with the observation that *GHRd3* is potentially fixed in ancient Eurasian hominins (i.e., Neanderthals and Denisovans), who may have been exposed to even greater seasonal environmental stress than most African hominins (*24*). Furthermore, the marked reduction in *GHRd3* allele frequencies coincides with the emergence of technologically advanced material culture, such as bone tools, fish hooks, and composite weapons, along with a concurrent population expansion, seemingly occurring independently in different parts of the world between 90 to 30 thousand years ago (*25*). Emerging technological innovations that may have allowed these populations to adapt to diverse environments (*26*) may also have acted as a buffer against the effects of fluctuating resource levels, thereby relaxing the selective pressures on the *GHR* locus. It is also noteworthy that the reduction in *GHRd3* allele frequency in Asia coincides with the Last Glacial Maximum (*27*). Thus, the combined effect of cultural and climate change could explain the rapid adaptive decrease in *GHRd3* allele frequencies in all human populations, the most dramatic of which was observed in Eurasia.

Based on our evolutionary model, we expected *GHRd3* to have pronounced phenotypic effects when resources are limited. Further, given the previous data suggesting sex-specific effects of *GHRd3*, we expected the phenotypic effects of *GHRd3* to be confined to males. To test these expectations and further investigate the biological effects of *GHRd3,* we developed a mouse model by generating a ∼2.5 kb deletion using CRISPR/Cas9. This deletion mimicked the human variant and removed the otherwise conserved exon 3 ortholog from the C57BL/6 genome. This procedure involved editing the genome of mouse embryos using a pair of single guide RNAs (sgRNAs) that complement the sequences flanking the third exon of the growth hormone receptor gene (*Ghr*) in the mouse genome (**Fig. 2A, Supplementary Methods**). The resulting founder mouse was a heterozygous (wt/d3) male, which was then used to generate offspring with wild-type (wt/wt) mice. We backcrossed the initial wt/d3 mice with wt/wt mice for at least 5 generations before conducting the experiments to prevent off-target effects. No deviations from Mendelian expectations were observed in the initial colony or subsequently established colonies of this mouse line. Further, we sequenced the genomes of 3 *Ghrd3* homozygous mice and found no evidence for off-target effects (**Supplementary Methods**). We confirmed mouse genotypes using both gel electrophoresis (**Fig. S8**) and expression analyses of exon 3 in the *Ghrd3* mice (**Fig. S9, Supplementary Methods**). To test the specific expectations outlined above, we established eight cohorts of 5 mice (**Table S2**) based on all possible combinations of sex (female or male), diet (constantly available, *i.e.*, *ad libitum (*AL*) or* 40% calorie restricted (CR)), and genotype (wt/wt or d3/d3) under controlled settings (**Fig. 2B**). We raised all of the cohorts under identical conditions until weaning at 30 days old and provided the differential diets for the next 30 days.

**Figure 2.**
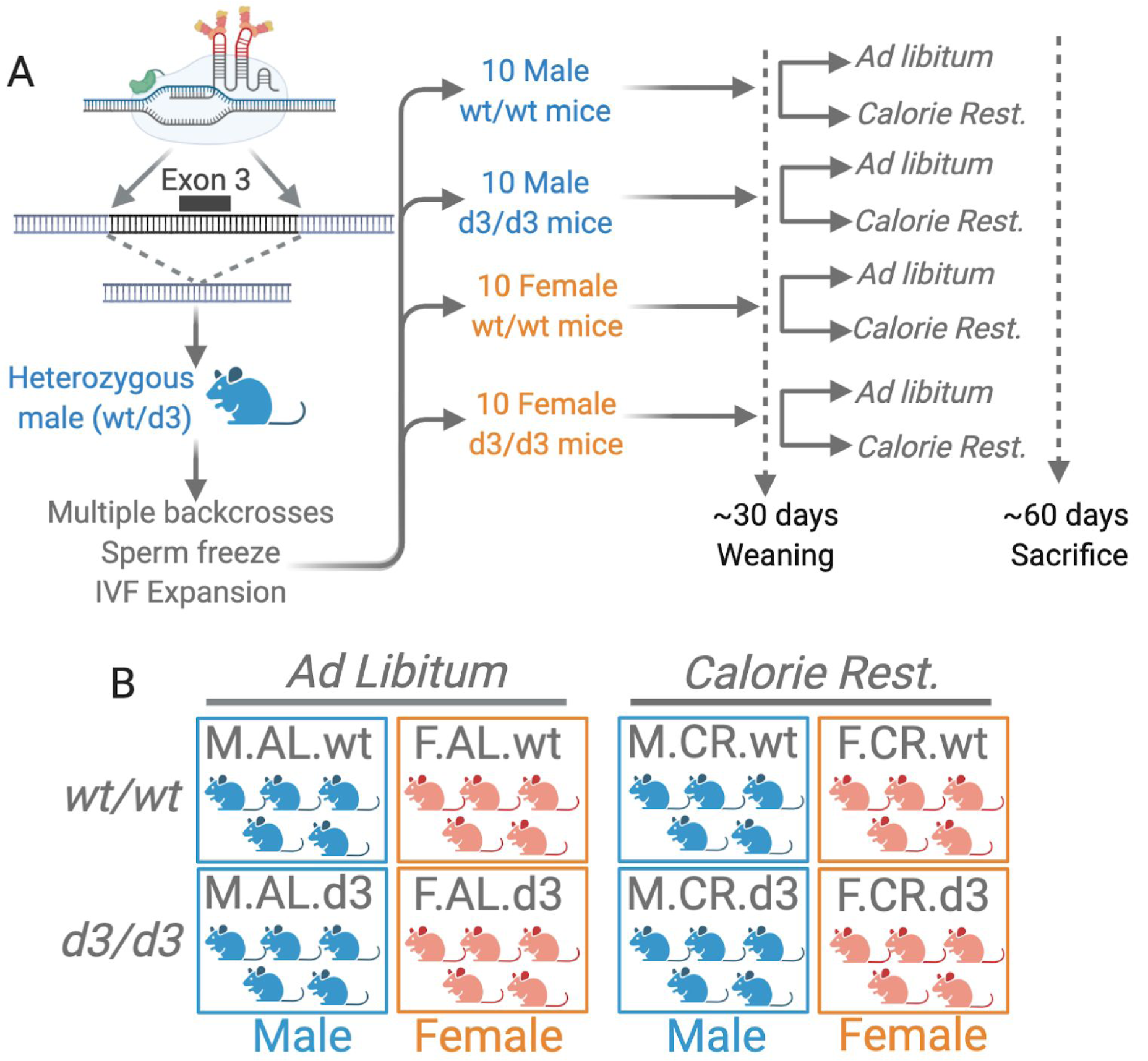
Experimental design. **A.** Generation of the mouse model using a CRISPR/Cas9 approach with sgRNAs targeting each side of the deletion. After multiple backcrosses, we raised the male and female mice with or without the deletion under *ad libitum* (*AL*) and 40% calorie-restricted (CR) dietary conditions. **B.** Based on the experimental design, we utilized 8 different cohorts, each having 5 mice.

At ∼60 days, we observed no significant effect of *Ghrd3* on the weight of mice that were fed the AL diet (**Fig. 3A, Table S2**). However, under CR, we observed that *Ghrd3* leads to a -8.4% (p=0.016, MW test) and a +4.75% change in mean weight in males and females, respectively. As a result, sexual differentiation in weight completely disappears under CR among d3/d3 mice. This is a remarkable finding considering that male mice are ∼39% and 20% larger in wt/wt mouse cohorts grown under AL and CR, respectively (p<0.01, MW test). Overall, our results are consistent with the notion that the phenotypic effect of *Ghrd3* is strong, but manifests in a sex- and environment-dependent manner.

**Figure 3.**
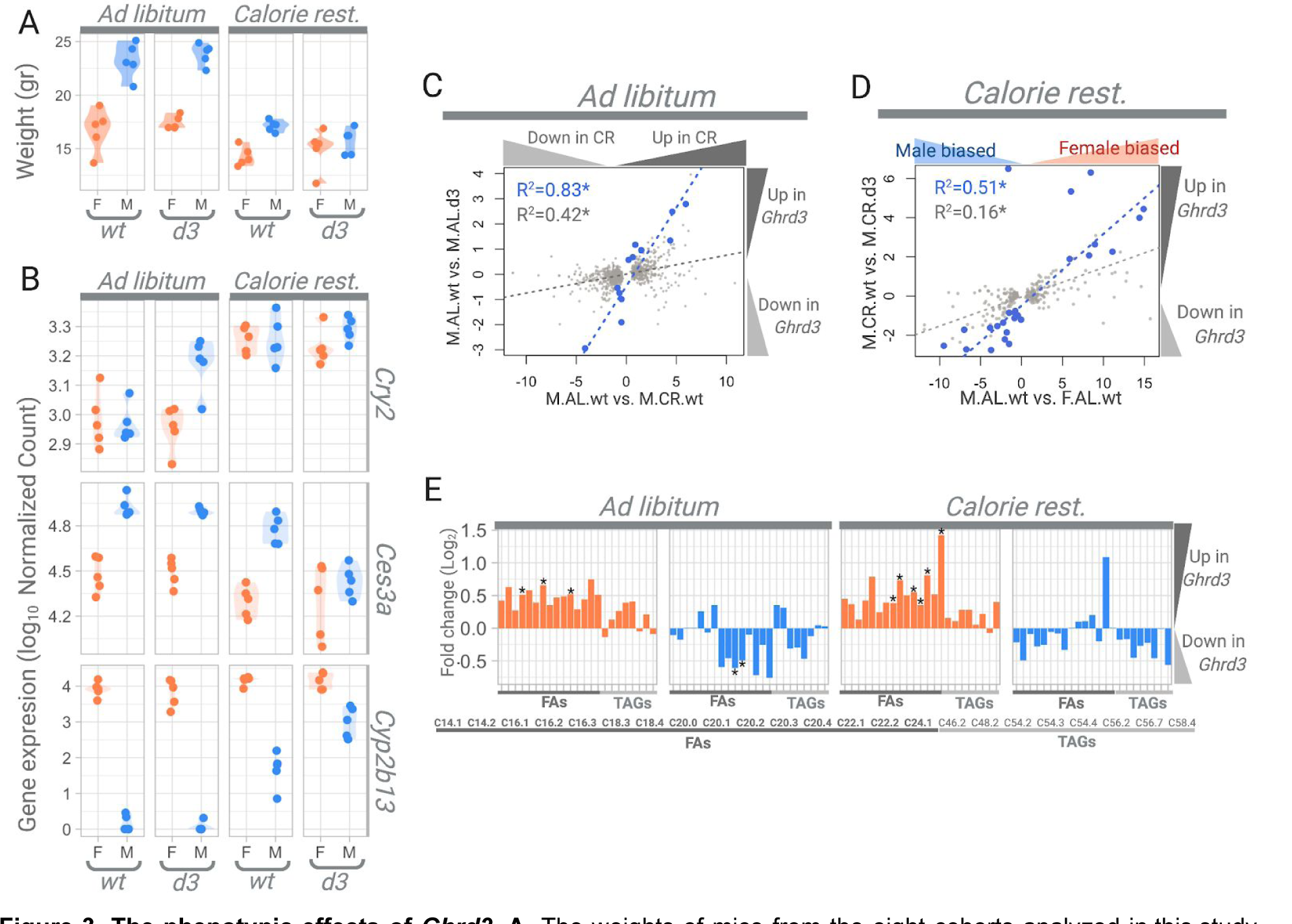
The phenotypic effects of *Ghrd3.* **A.** The weights of mice from the eight cohorts analyzed in this study. For the entire figure, female and male mice are labeled by orange and blue, respectively. **B.** Examples of specific genes that show significant differences between wt/wt and d3/d3 male mice. *Cry2* is an example of a circadian gene that shows a significant difference in expression between d3/d3 and wt/wt mice given an *AL* diet. *Ces3a* is an example of a gene with male-biased expression under normal conditions but is significantly reduced in expression in male d3/d3 mice under a CR diet. *Cyp2b13* is an example of a gene with female-biased expression under normal conditions but significantly increased expression in male d3/d3 mice under calorie restriction. These two genes with sex-biased expression exemplify the female-like expression of the d3/d3 male mouse livers under calorie restriction. **C.** The correlation between the effect of calorie restriction (x-axis) and *Ghrd3* under an *AL* diet (y-axis). Specifically, the x-axis shows the effect size of calorie restriction by log_2_ ratios of gene expression levels between wild-type male mice that were fed *AL* and CR diets. The Y-axis shows the log_2_ ratio of gene expression levels between wt/wt and d3/d3 male mice that were grown under *AL* diets. The light gray dots show the genes that have significantly (p_adj_<0.01) different expression levels between wild-type male mice that were fed *AL* and CR diets. The blue dots show the genes that have significantly (p_adj_<0.01) different expression levels between wt/wt and d3/d3 male mice that were grown under *AL* diets. The color-matched regression lines and R^2^ values are provided by these two distributions. **D.** The correlation between the effect of sex (x-axis) and *Ghrd3* under calorie restriction (y-axis). Specifically, the x-axis shows the log_2_ ratio of gene expression levels between wild-type male and female mice that were fed an *AL* diet. The Y-axis shows the log_2_ ratios of gene expression levels between wt/wt and d3/d3 male mice that were given a CR diet. The lighter blue dots show the genes that have significantly (p_adj_<0.01) different expression levels between males and females, while darker blue dots show the genes that have significantly (p_adj_<0.01) different expression levels between wt/wt and d3/d3 male mice that were grown under a CR diet. The color-matched regression lines and R^2^ values are provided by these two distributions. **E.** Log_2_ fold changes between wt/wt and d3/d3 mice, showing sex-specific impacts of *Ghrd3* on the levels of FAs (fatty acids) and TAGs (triacyglycerides) in the blood serum of females and males. The significant changes (nominal p<0.05, t-test) are marked by *.

To understand the mechanistic underpinnings of the biological effects of *Ghrd3*, we conducted a comparative transcriptomics analysis of the liver tissues of mice from the eight cohorts at ∼60 days (**Table S3**). At this age range, the mice have reached a point of sexual maturity but were still in a period of growth and development (*28*). The liver is a natural tissue choice for this analysis because *GHR* is expressed abundantly in the liver and its specific functions in this tissue have been extensively studied (*29*). Moreover, the overall expression trends of mouse livers for both sexes are well studied (*30*). Since growth hormone (GH) secretion patterns are circadian and sex-specific (*31, 32*), we expected the downstream effects of *Ghrd3* to be highly variable because they are intrinsically dependent on GH availability. Thus, only the strongest effects would likely be significant when comparing wt/wt and d3/d3 liver transcriptomes. Indeed, we found no genes with significant expression differences when we compared F.AL.wt and F.AL.d3 or F.CR.wt and F.CR.d3 cohorts (see **Figure 2B** for cohort descriptions). These results are consistent with the notion that *Ghrd3* has a minimal effect in females, independent of diet, at least at the two-month developmental snapshot. In contrast, we found 13 differentially expressed genes when we compared M.AL.wt and M.AL.d3, and 28 differentially expressed genes when we compared M.CR.wt and M.CR.d3 cohorts (**Table 1**, p_adj_<0.01, Wald test with Benjamini-Hochberg multiple testing correction). Surprisingly, these sets of genes do not overlap, indicating that the effect of *Ghrd3* on gene expression in male mouse livers differs based on dietary conditions.

To further understand this phenomenon, we first conducted an enrichment analysis of functional categories for the genes that are differentially expressed between M.AL.wt and M.AL.d3 livers (**Table S4, Supplementary Methods**). We found that 5 out of the 13 genes (∼38%) are related to circadian rhythm processes, indicating a significant enrichment from stochastic expectations (FDR=2.33x10^-6^). Notably, these include the upregulation of three primary circadian rhythm regulators, *Per3*, *Cry2*, and *Ciart* (**Table 1**). A closer inspection of the expression trends in all the mouse cohorts revealed that the same circadian rhythm genes are also upregulated in wild-type mice as a response to calorie restriction (e.g., *Cry2*, **Fig. 3B**). Indeed, circadian response to dietary change directly interacts with the GH pathway and involves the regulation of *Per* and *Cry2* expression (*33, 34*). The expression levels of circadian rhythm regulators in M.AL.d3 mice resemble those of wild-type mice under CR.

Calorie restriction dampens growth hormone cyclicity in mice, leading to the flattened pulsation of its downstream signal (*35*). This flattening is partially compensated by the activation of the circadian rhythm pathway, involving *Per* and *Cry* (*33, 34*). Since *Ghrd3* transduces growth hormone signaling ∼30% faster than the full-length isoform (*8*), we hypothesize that it has a similar dampening effect, reducing the pulsating downstream signaling in the growth hormone pathway even under an AL diet. Supporting this hypothesis, we found a significant correlation between the gene expression response to calorie restriction in wild-type mice and the effect of *GHRd3.* Specifically, the expression change from M.AL.wt to M.AL.d3 significantly correlates with the expression change from M.AL.wt to M.CR.wt (p< 10^-5^, **Fig. 3C**). Overall, our results support the notion that the effect of *GHRd3* is similar to calorie restriction in male mice under normal dietary conditions.

Next, we conducted an enrichment analysis on the 28 genes that are differentially expressed in M.CR.d3 mice as compared to M.CR.wt mice. We found that 13 of these 28 genes are involved in “HNF4A-Dependent sex-specific differences in mice liver” (*36*), and were highly enriched (∼46%, FDR=1.62X10^-12^). Indeed, a closer inspection revealed that multiple well-established *male-biased* genes (e.g., major urinary protein (*Mup)* genes and carboxylesterases (Ces3a and 3b)) were downregulated in M.CR.d3 livers when compared to M.CR.wt livers, while notable *female-biased* genes (e.g, *Cypb13*, *Cyp2a4*, *Sult3a1*, *Fmo3*) were upregulated (**Table 1**). When we investigated the expression of those genes in all 8 cohorts, we found a remarkable similarity between the expression of these genes in M.CR.d3 mice and female mice in general, regardless of the diet and genotype (e.g., *Cyp2b13* and *Ces3a;* **Fig. 3B**). In that regard, the expression of these genes mimics the weight variation. To recap, M.CR.d3 mice are significantly lighter than M.CR.wt mice, making them virtually identical in weight to F.CR.d3 mice (**Fig. 3A**).

Based on these observations, we hypothesize that *Ghrd3* leads to the female-like gene expression in male livers under calorie restriction, which potentially leads to the observed size reduction. It is important to note here that all of these genes were shown to be downstream gene targets of the *Hnf4α* and *Stat5b* pathways and that their concerted signaling regulates several sexually dimorphic genes in the liver (*37*). We also found a significant correlation between the effect of *Ghrd3* in males under calorie restriction (measured by comparing M.CR.wt and M.CR.d3 cohorts) and sex-specific differences in gene expression under AL dietary conditions (measured by comparing F.AL.wt and M.AL.wt cohorts) (**Fig. 3D**, p<10^-4^).

These results are consistent with a model where *Ghrd3* leads to a female-like liver expression in male mice under calorie restriction. It was suggested that the pulsatile secretion of growth hormone promotes male-biased gene-expression while suppressing female-biased gene expression (*37*). Conversely, persistent growth hormone treatment disrupts the intermittent nature of Stat/Jak activity, which leads to the promotion of female-biased gene expression while suppressing male-biased expression. As mentioned before, *GHRd3* was shown to transduce growth hormone signaling faster than the ancestral type of the protein (*8*). It is plausible that this stronger binding leads to a more persistent downstream signaling. This effect is likely compensated under an *AL* diet through the upregulation of the expression of circadian rhythm genes as described above. However, this compensation mechanism may not be strong enough to overcome the combined dampening effect of *Ghrd3* and calorie restriction. Therefore, we predict that *Ghrd3* under calorie restriction dampens the growth hormone signaling cyclicity, leading to the female-like gene expression in the male liver. These results also suggest that the smaller size observed in male d3/d3 mice under calorie restriction is at least partially the result of these transcriptome-level effects we document in the male liver.

Given previous reports on the effect of *GHR* on lipid metabolism (*29*), as well as our own results that identify multiple lipid metabolism-related genes (e.g., multiple *Mup* and *Ces* genes), we investigated the downstream effects of *Ghrd3* on the blood serum lipid composition using liquid chromatography-mass spectrometry (**Table S5**). We chose to focus on serum lipids because we wanted to understand the global metabolic effect of *Ghrd3* at the organismal level. Furthermore, serum lipids are routinely measured for diagnostic purposes, and we wanted to construct a comparable dataset for future studies. We documented a striking, opposite effect of *Ghrd3* on fatty acid and triglyceride composition in the serum of males and females, independent of diet (**Fig 3E**). The opposing effect is particularly prominent for fatty acids. Specifically, we found that all 15 fatty acid species that we analyzed were upregulated in female *Ghrd3* mice compared to their wild type counterparts, whereas the majority of these lipids were downregulated in male *Ghrd3* mice. The effects of *Ghrd3* on serum lipids seem to be similar in mice that are fed with AL and CR diets and thus differ from the observations in the liver. This observation is consistent with the previously proposed notion that the effect of the growth hormone receptor varies between different tissues (*38*).

Our results provide one of the very few human examples (*39*) where the effects of a common genetic variant are sex- and environment-dependent. In that regard, we suggest that *GHRd3* has important ramifications for metabolic disorders, such as obesity and diabetes, but only for males within particular environmental contexts. For example, it was reported that *GHRd3* has a preventative impact on type 2 diabetes. However, in the small number of diabetes patients who are homozygous for *GHRd3*, a significant increase in further metabolic complications was observed (*17*). This parallels our observation that a significant size difference was observed between wt/wt and d3/d3 mice only under calorie restriction for males. Similarly, we expect the reported effects on birth and placental size, time to menarche, and longevity to vary significantly for different dietary and endocrinological contexts. Following this thread, we expect that *GHRd3* may have other important roles that would be visible only when specific sexes, environmental conditions, and developmental stages are investigated. Our study has characterized the effect of homozygous *Ghrd3.* Given that this gene codes for a protein that self-dimerizes for it to function properly, the functional impact of *Ghrd3* in heterozygous individuals remains a fascinating question. Our mouse model makes it possible to test all these different perspectives in future studies.

The sex-specific effects of *GHRd3* raise several evolutionary considerations, especially because the growth hormone pathway is a major driver of sexual dimorphism in mammals (*40*). Moreover, multiple genes in this pathway were recently implicated in sexually antagonistic transmission distortions in humans (*41*). In this regard, *GHRd3* provides an interesting case.

The traditional understanding of the evolution of sex-specific effects of genetic variation first considers sexual conflict where the functional effect of the variant leads to increased fitness in one sex but decreased fitness in the other (*42*). This conflict leads to maintenance of the variation in the population through balancing selection until the conflict is resolved by additional genetic variation that moderates the effect of the variant in a sex-specific manner. *GHRd3* does not fit this scenario. Instead, the sex-specific effect of *GHRd3* is instantaneous because the growth hormone pathway is already operating in a sex-specific manner. This observation raises questions about the extent of the evolutionary effects of genetic variation in pathways that already operate in a sex-specific manner.

Our insights into this genetic variation at the *GHR* locus bring forth several questions concerning recent human evolution. The initial allele frequency increase of *GHRd3* across evolutionary time coincides with unstable environments that mark recent hominin evolution (*43*). Given its enhanced effect under calorie restriction, it is plausible that *GHRd3* provided a fitness advantage to early hominins, perhaps facilitating adaptation to new environments during *homo* migrations out of Africa starting ∼1 million years ago. Indeed, our finding that *Ghrd3* is protective against the severe consequences of malnutrition in male children supports this hypothesis. The sex-specific nature of this adaptation may be better understood by considering times of nutritional stress and assuming that the increased size in males is a derived trait. During these periods, survival may be a stronger adaptive force than the fitness benefits of increased size, causing smaller sizes in males to be favored (*44*). Thus, *GHRd3* may be favored under environmental stress because it increases survival even though it reduces sexual dimorphism and thus reduces competitiveness for mate choice among males. The recent decrease in the allele frequency of *GHRd3* in humans, most notably in Eurasia, coincides with technological transitions, perhaps providing a buffer against cycles of famine. In such stable environments, the effect of sexual selection would be relatively higher, favoring the non-deleted ancestral allele.

## Data Availability

All the fastq files from the genome and RNAseq experiments are being uploaded to NCBI sequence read archive (https://www.ncbi.nlm.nih.gov/sra). All other data are available in the Supplementary Materials and Tables.

## Acknowledgments

Funding: This work was financially supported by the National Science Foundation (O.G., no. 1714867). Work in the Mu lab was also supported by grants from the BrightFocus Foundation (G2016024) and the National Eye Institute (R01EY020545 and R01EY029705). The mouse work in Jackson Laboratories was supported by an NIH grant (#P30AG038070). S.R. received funding during his first two years of research from the Collaborative Learning and Integrated Mentoring in the Biosciences (CLIMB) program. Human genetic studies of severe malnutrition were supported by the Doris Duke Charitable Foundation (#2013096) and the USDA, ARS cooperative agreement (58-3092-5-001). Author contributions: O.G. designed, oversaw and managed the study. He contributed to analysis in each step. X.M. helped design the experiments and oversaw the study. M.S. conducted transcriptomics and population genetics analysis. S.R. conducted the majority of the mouse work and genotyping.

A.J.P. and G.E.A-G conducted lipidomics analysis, S.N. and Y.S. conducted simulation-based selection analysis. F.M. helped in tissue collection in mice. L.R. and G.C. designed and oversaw mice experiments in Jackson Laboratories. N.A.B., N.J.H., and N.L. contributed analysis of malnutrition outcome association study. C.L. and Q.Z. contributed to transcriptomics and genomic sequencing for looking for potential off-target effects from CRISPR-cas9 experiments. O.G., S.R., and M.S. wrote the manuscript and prepared the figures. Competing interests: The authors declare no competing interests.

We are grateful to the Roswell Park Cancer Institute Gene Targeting and Transgenic Resource, especially Aimee Stablewski, for their help in establishing the *Ghrd3* mouse line. We are grateful for the help of George (PJ) Perry, and Audrey Arner regarding the navigation of the UK Biobank dataset. We would like to acknowledge Kirsten Dean and Victoria Gellatly for their help in genotyping. We thank our colleagues Drs. Xu-Friedman, Taylor, Lynch, Halfon, and Albert for their constructive criticism throughout this study.

## SUPPLEMENTARY MATERIALS

### Supplementary figures

**Figure S1.**
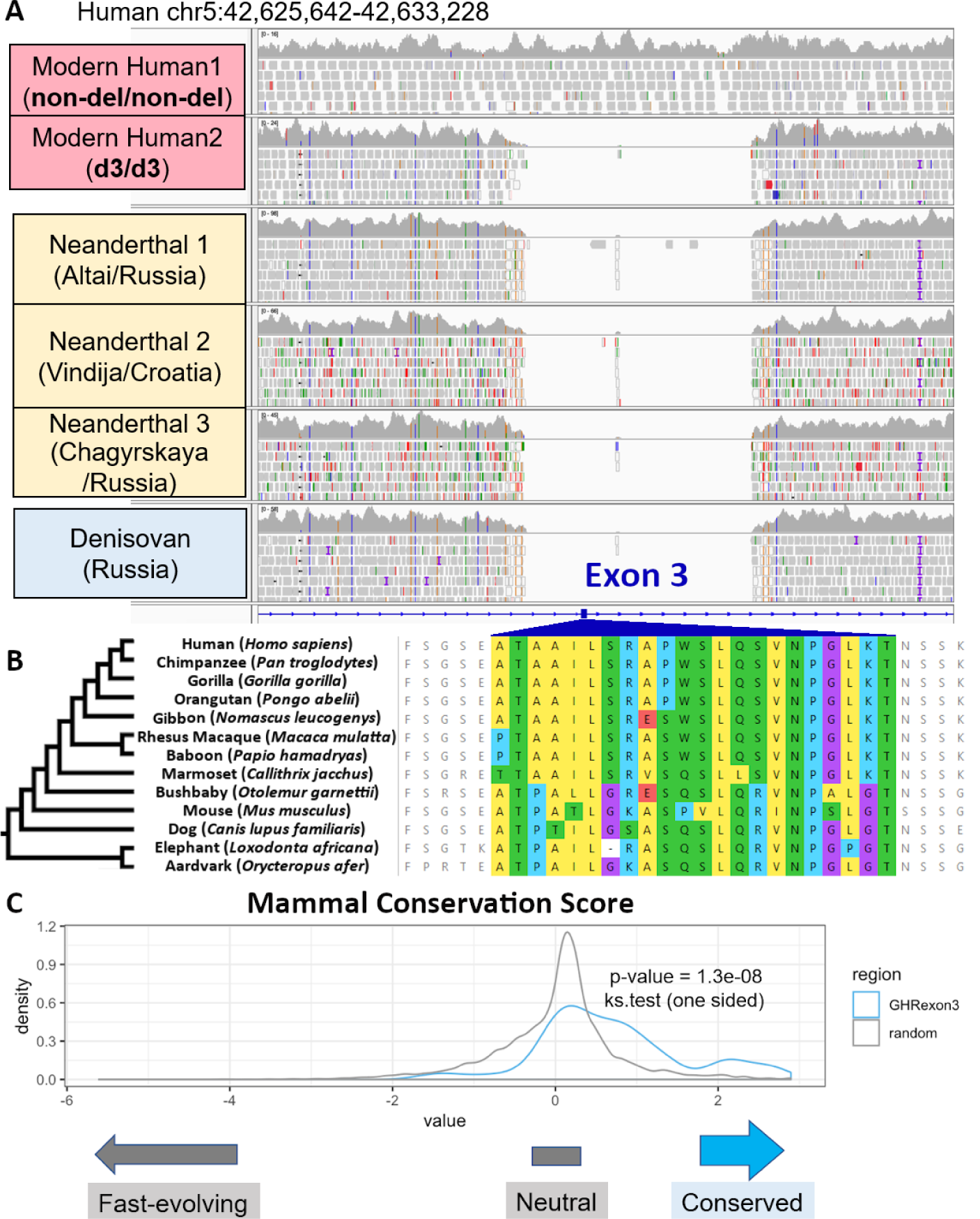
**A.** *GHRd3* in modern and ancient hominin genomes. These browser snapshots show the genome assembly (Hg19) of a human with the ancestral, homozygous non-deleted genotype and another with a homozygous deleted genotype that shows no reads mapping to the deletion region (top two rows). Similarly, sequences from 3 Neanderthal and 1 Denisovan assemblies were mapped to this region and show a clear signature of the deletion with breakpoints indistinguishable from the deletion observed in modern humans. **B.** The coding sequence alignment of exon 3 among mammals, which indicates near-complete preservation of the amino-acid sequence in primates and clear conservation across mammals. **C.** A conservation score (phastCons46.mammals in UCSC Genome Browser) comparison of *GHR* exon 3 sequences with 20,000 randomly chosen sites from chromosome 5.

**Figure S2.**
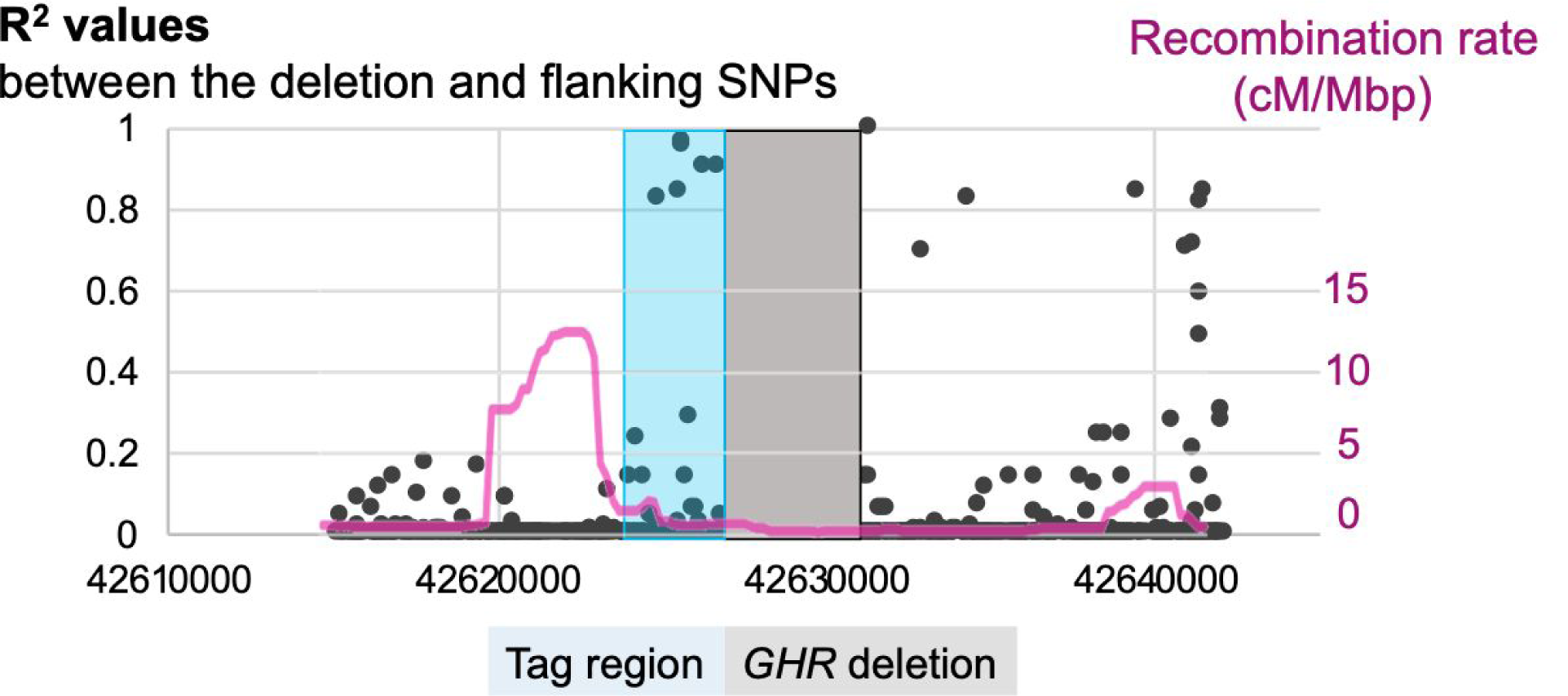
Linkage disequilibrium between SNPs near *GHRd3* and the deletion. This figure shows R^2^ values calculated for the *GHR* deletion and its neighboring SNPs (black dots, left scale), along with recombination rates for the locus (the pink line, right scale). Recombination rates were retrieved from the 1000 genomes selection browser (http://hsb.upf.edu/). Based on the linkage disequilibrium, we selected Hg19: chr5: 42624748-42628325 as the tag region.

**Figure S3.**
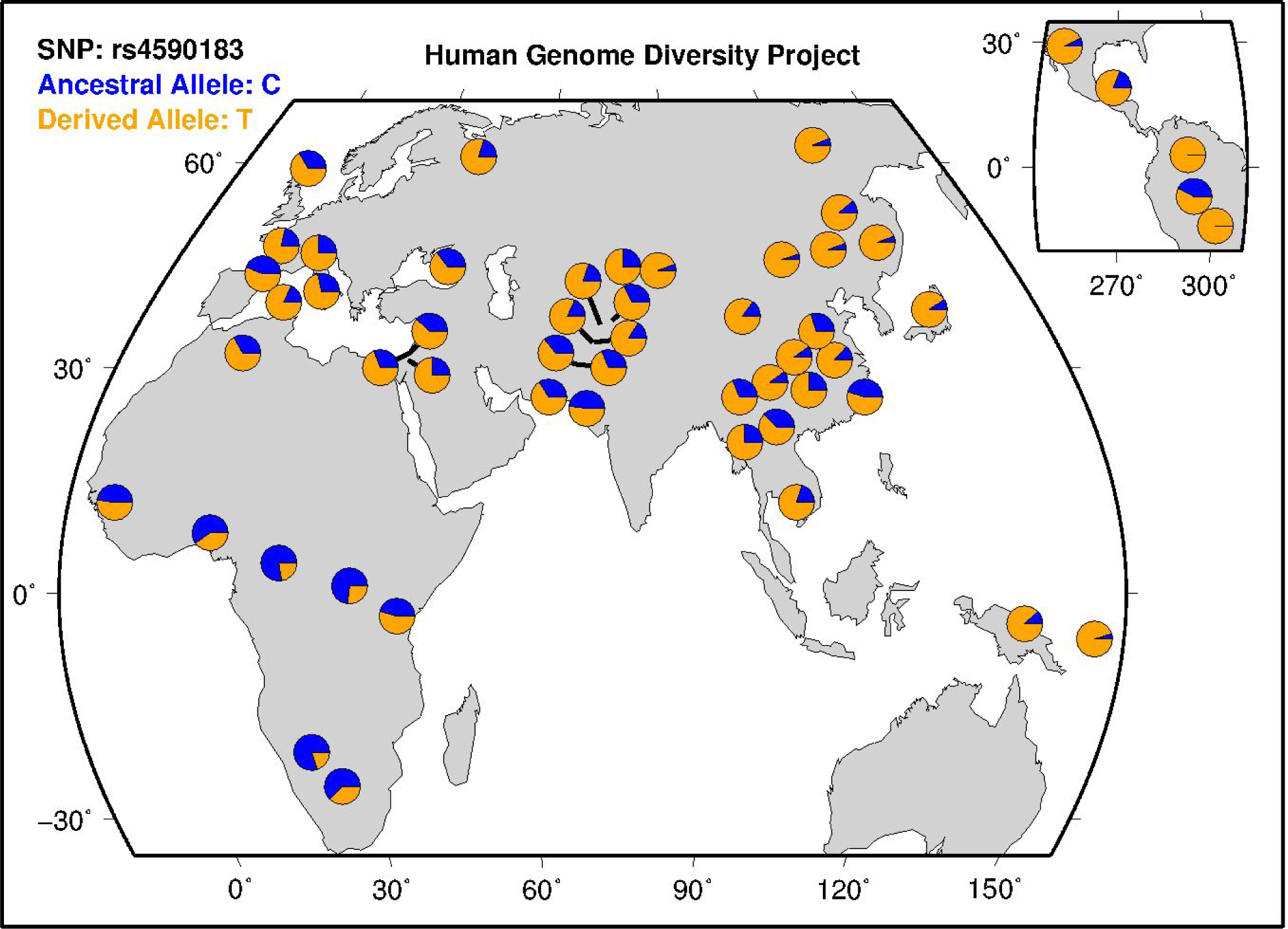
*GHRd3* tag SNP frequency map. Global frequencies of the *GHRd3* tag SNP from the Human Genome Diversity Project (https://www.hagsc.org/hgdp/); The C allele tags the derived deletion, despite being the ancestral allele.

**Figure S4.**
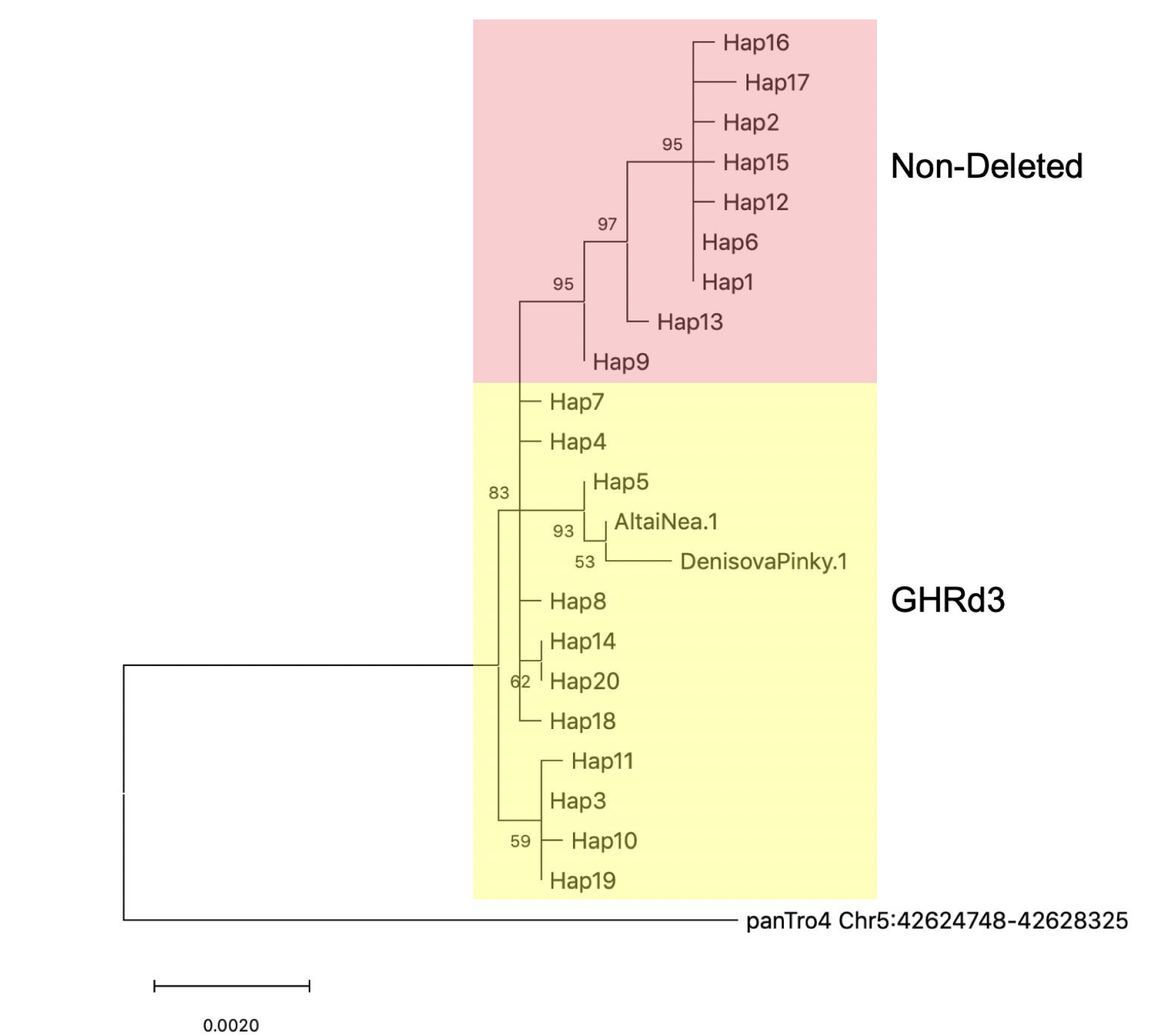
*GHR* exon 3 phylogenetic tree. This phylogenetic tree was constructed using randomly selected haplotypes that harbor the deleted and non-deleted *GHR* exon 3 alleles. It is clear from this phylogeny that haplotypes harboring the deleted allele are more diverse and coalesce earlier than those that harbor the non-deleted allele. It is also noteworthy that both the Altai Neanderthal and Denisova genomes cluster with the haplotypes harboring the deleted allele.

**Figure S5.**
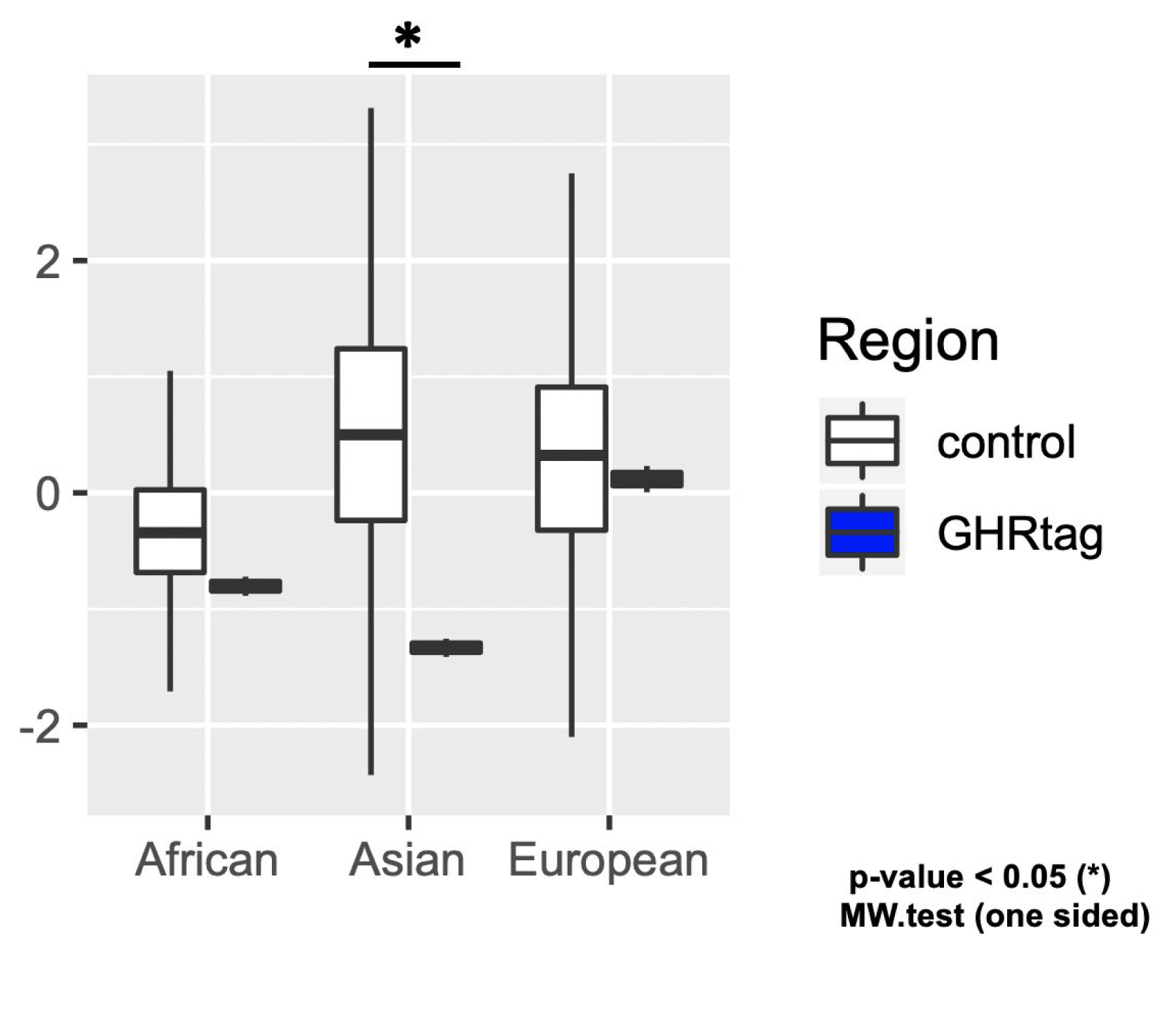
Tajima’s D values for all haplotypes. Tajima’s D values between the *GHRd3* tag region (Hg19: chr5: 42624748-42628325, blue) and 500 randomly selected regions across the genome (white) for three populations.

**Figure S6.**
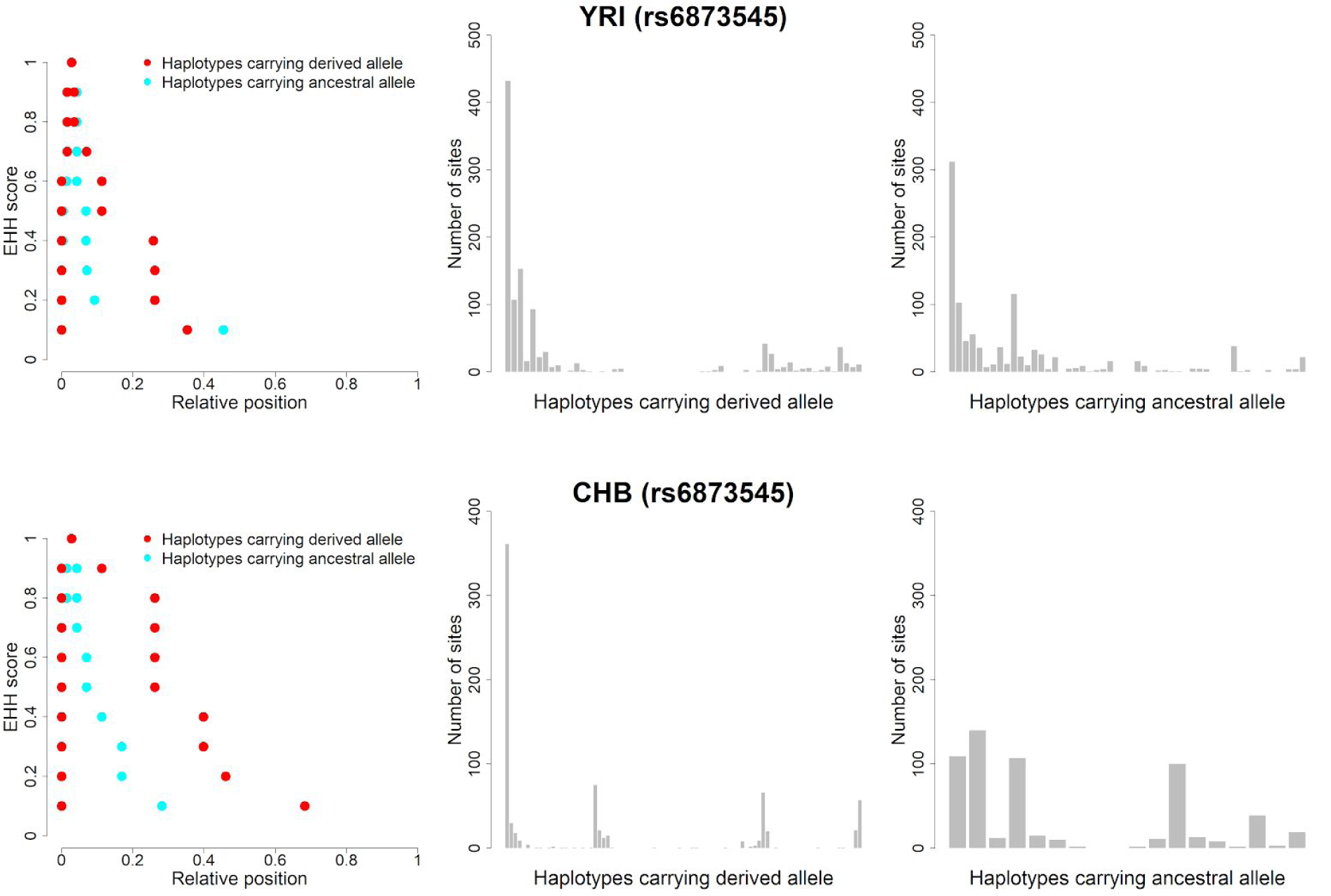
Approximate Bayesian computation testing different models. Decays of extended haplotype homozygosity (EHH) (left) and site frequency spectrum (SFS) (middle for the derived allele and right for the ancestral allele) in YRI (top) and CHB (bottom) populations. The EHH shows the probability that two randomly chosen haplotypes are homozygous at all SNPs within a given distance from a focal SNP site, which in this case is a tag variant for *GHRd3* (rs6873545). This measure is depicted as a value between 0 and 1. To fully capture the haplotype structure in this region, we recorded the physical positions of the variable sites on each side of the focal SNP where the EHH value decreased from 0.9 to 0.1, in steps of 0.1, for derived (red dots) and ancestral alleles (light blue dots) separately. The SFS covers the entire range of allele frequencies from singletons to mutations shared across all chromosomes, providing the local reduction in nucleotide diversity and the distortion of the SFS in a population. There were 113 chromosomes carrying the derived alleles and 103 carrying the ancestral alleles in the YRI samples; For the CHB samples, these numbers were 170 and 36, respectively. The bin size was set as “2”. Therefore, the total number of bins for those chromosomes was 57 and 52 for YRI and 85 and 18 for CHB. We used all of the data shown in these plots as a set of summary statistics for the analysis of approximate Bayesian computation (ABC) (see Supplementary information).

**Figure S7.**
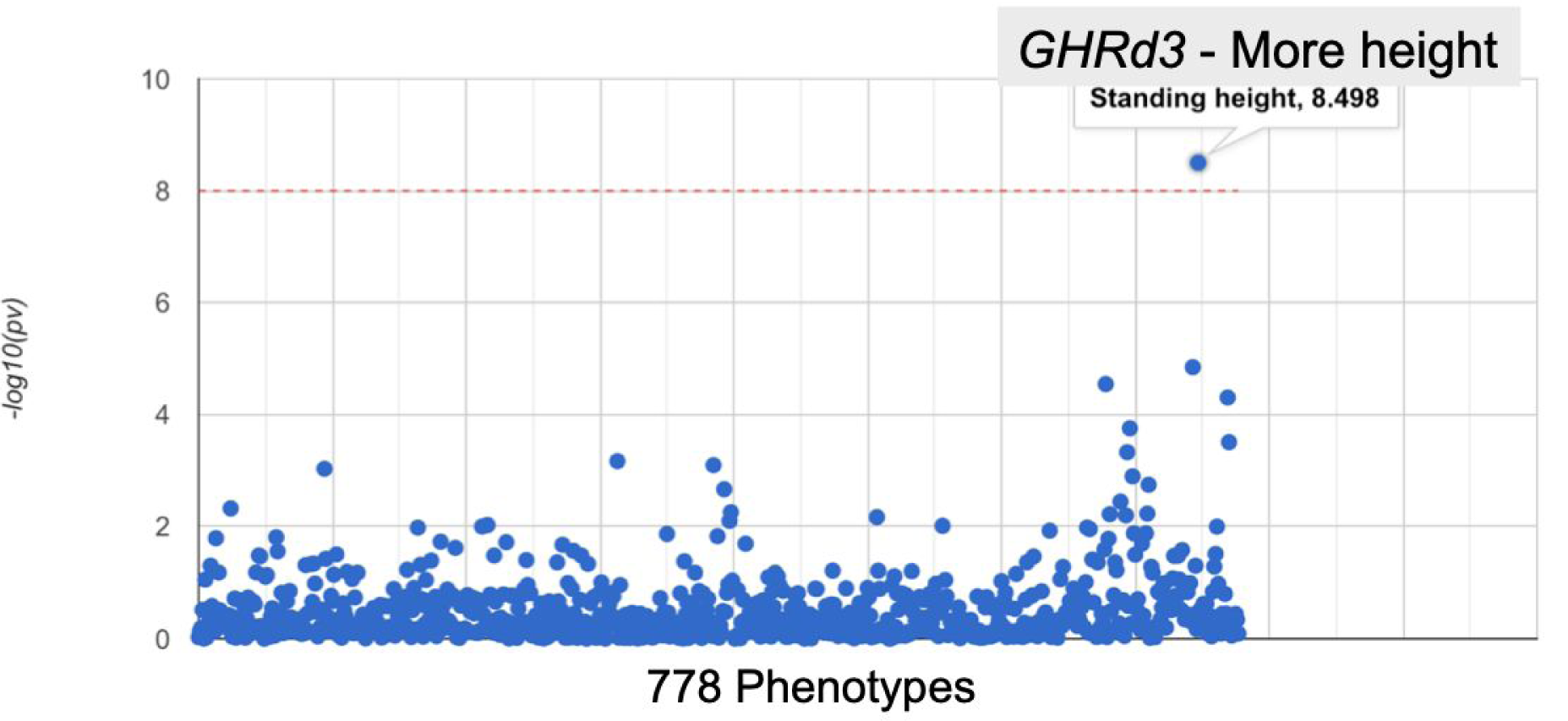
Phewas analysis on the *GHRd3* tag SNP. The UK biobank PheWas (http://geneatlas.roslin.ed.ac.uk/phewas/) results using the tag SNP (rs4073476).

**Figure S8.**
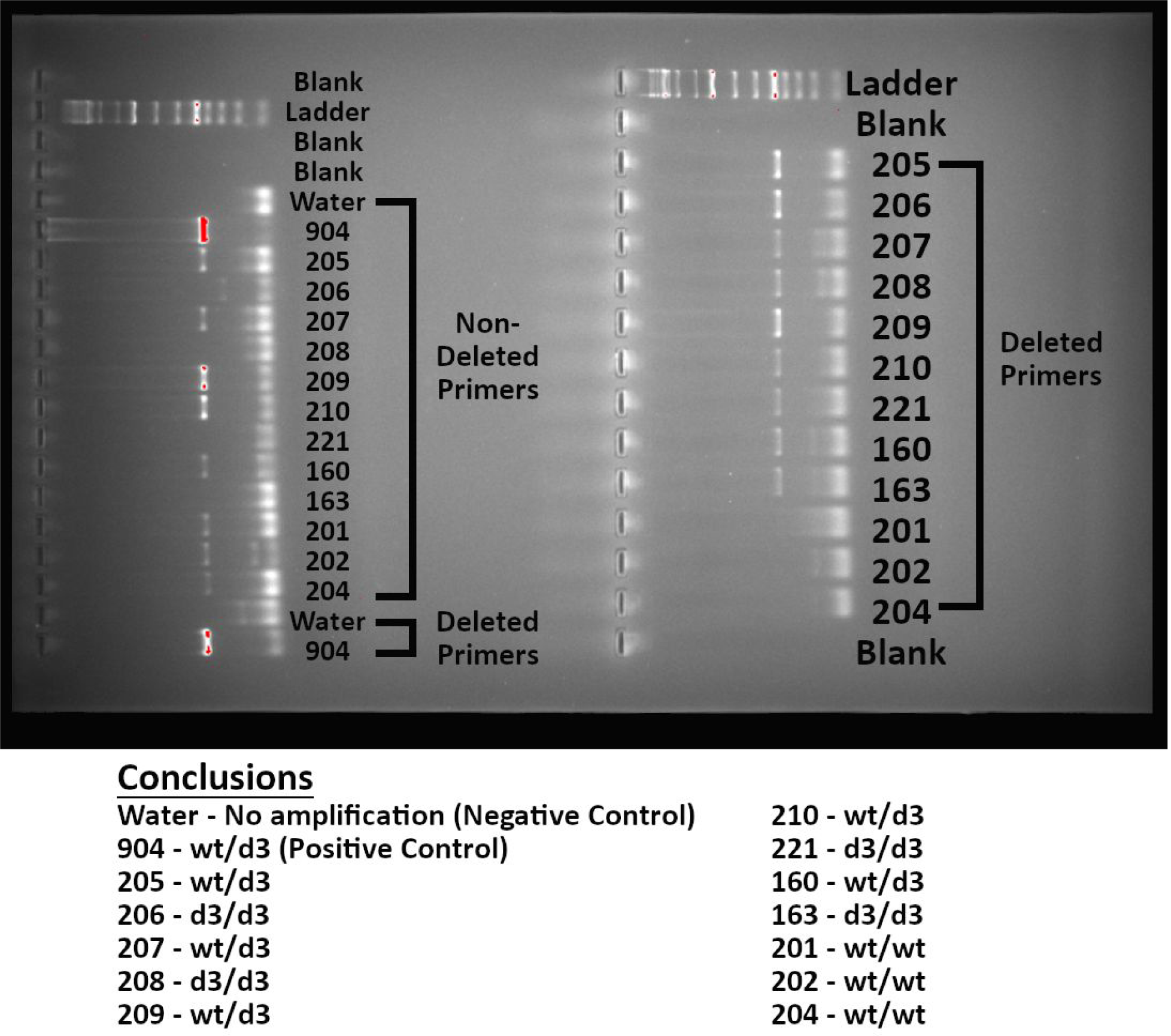
A gel exemplifying our genotyping approach. A two percent agarose gel showing polymerase chain reaction products. Two sets of primers were used. A sample that showed amplification with the non-deleted primers, but not the deleted primers would be deemed wt/wt; A sample that showed amplification with the deleted primers, but not the non-deleted primers would be deemed d3/d3; A sample that showed amplification with both sets of primers would be deemed wt/d3. Water was used as a negative control and a known heterozygote (904) was used as a positive control.

**Figure S9.**
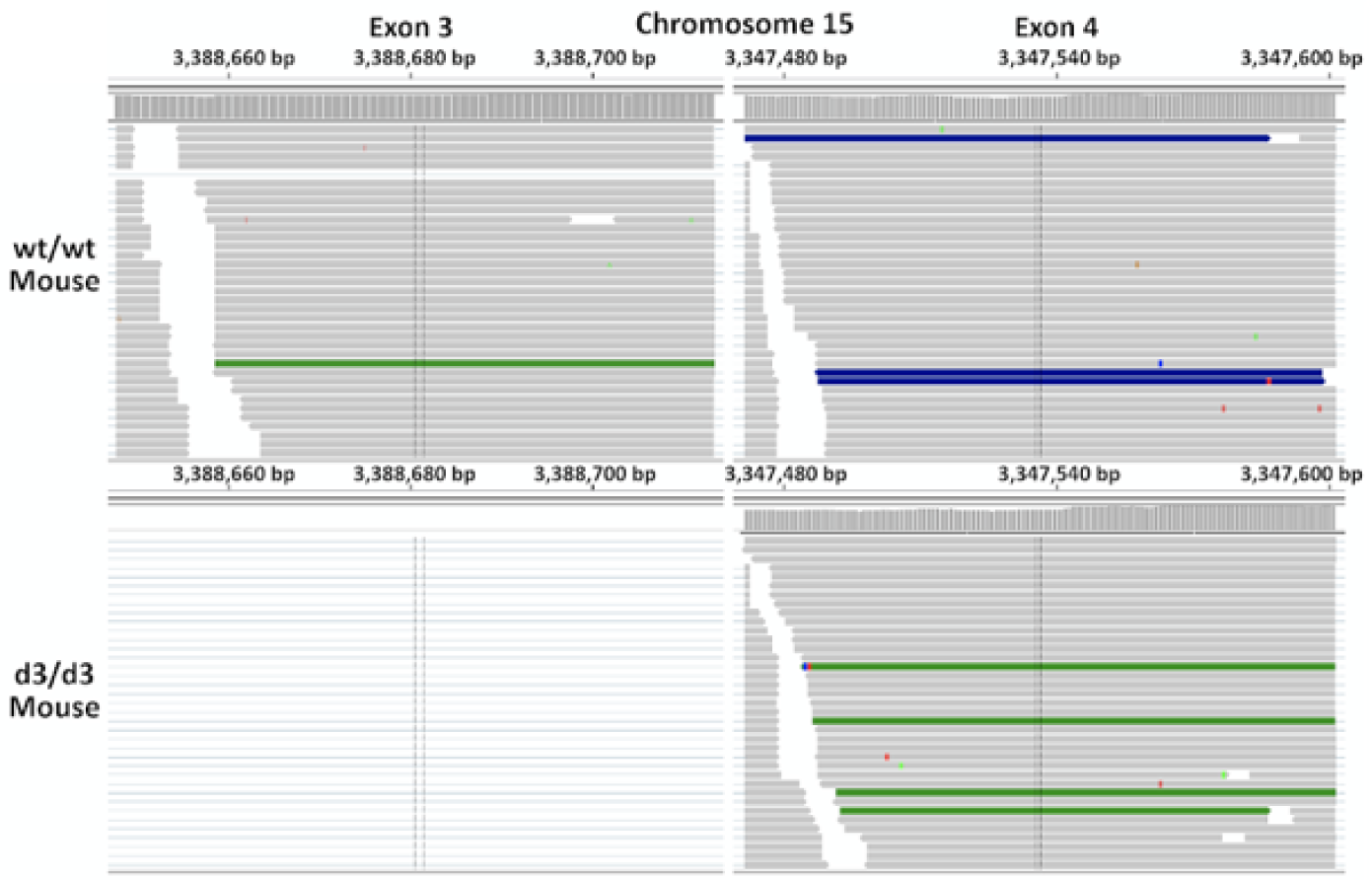
RNAseq exonic read-depth from a wt/wt and d3/d3 mouse for exons 3 and 4. Each horizontal grey line represents a single read. The d3/d3 mouse expresses *GHR* exon 4 in similar levels as compared to the wt/wt mouse, but does not express exon 3.

### SUPPLEMENTARY METHODS

#### Haplotype network and read-depth analysis

We used the program vcftools (0.1.16) (*45*) to calculate the R^2^ values between *GHRd3* (esv3604875) and flanking SNPs to set a target region (Hg19: chr5: 42624748-42628325). A vcf file of the target region obtained from the 1000 Genome Project phase 3 dataset (*46*), the hg19 reference genome, the chimpanzee reference genome (*47*), the Altai Neanderthal genome, and the Denisovan genome (*48, 49*) were all used by the program VCTtoTree (V3.0.0) (*50*) to draw the haplotype networks. PopART (Version 1.7) (*51*) was used for the visualization of the network (**Fig. 1B**). The program rworldmap (*52*) was used to visualize the global allele frequency data from the 1000 Genome Project phase 3 dataset (*46*).

#### Conservation Analysis

We obtained the *GHR* coding sequence alignment from **Fig. S1B** through the UCSC genome browser by utilizing its “Other Species Alignments” function (MAF table: multiz100way) (*53*). This alignment was then viewed and exon 3 was highlighted using the program Molecular Evolutionary Genetics Analysis (MEGA v 10.0.5) (*54*). The software program bedtools (v2.27.1) (*55*) was used to obtain 20,000 random sites from chromosome 5, where the *GHR* gene is located, for the conservation analysis. The phastCons46way.placental datasets (*56*) from the UCSC Genome Browser were used to compare the conservation scores of *GHR* exon 3 and the randomly selected regions (**Fig. S1C**).

#### Neutrality Tests

To obtain random single nucleotide variants, we used bedtools (v2.27.1) (*55*) to construct random chromosomal coordinates on chromosome 5. We used 500 random regions for the *GHRd3* locus comparison. Tajima’s D (*57*) values for merged haplotypes (**Fig 1B** and **Fig S5**) and XP-EHH (**Fig. 1C**) (*13*) values were downloaded from the 1000 Genomes selection browser (*58*) and used for the bins containing the target region (Hg19: chr5: 42624748-42628325) and control region. To increase sample numbers for our Tajima’s D calculations we extended the analysis beyond the CEU, YRI, and CHB populations. The European category was expanded to include the following populations: Utah residents with Northern and Western European ancestry (CEU), Toscani in Italy (TSI), Finnish in Finland (FIN), British in England and Scotland (GBR), and Iberian populations in Spain (IBS); The African category was expanded to include the following populations: Gambian in Western Division in the Gambia (GWD), Mende in Sierra Leone (MSL), Esan in Nigeria (ESN), Yoruba in Ibadan, Nigeria (YRI), and Luhya in Webuye, Kenya (LWK); The east Asian category was expanded to include the following populations: Han Chinese in Beijing, China (CHB), Japanese in Tokyo, Japan (JPT), Southern Han Chinese, China (CHS), Chinese Dai in Xishuangbanna, China (CDX), and Kinh in Ho Chi Minh City, Vietnam (KHV). The visualization was constructed through ggplot2 (*59*).

For the analysis of LD patterns in different site frequency spectra, we followed the approach outlined by Fujito et al. (*14*). We used the two-dimensional site frequency spectrum (*14, 60*) to detect the signature of selection on *GHRd3*. This method can eliminate the effect of recombinations, which affect haplotype structures. We used the 1000 Genome meta-populations (Africa, East Asia, and Europe) for this analysis. By combining the site frequency spectrum method and coalescence simulations, we detected the signature of selection on the *GHR* non-deletion allele (*p* < 0.05) with the estimated date of selection onset being 27.5ky. The results of this analysis are summarized in the table below.

|  | Africa:n=1322 |  |  |  | East Asian:n=1008 |  |  |  | European:n=1006 |  |  |  |
| --- | --- | --- | --- | --- | --- | --- | --- | --- | --- | --- | --- | --- |
| ID | esv3604875 |  |  |  | esv3604875 |  |  |  | esv3604875 |  |  |  |
| position | 42628418 |  |  |  | 42628418 |  |  |  | 42628418 |  |  |  |
|  | der. |  | anc. |  | der. |  | anc. |  | der. |  | anc. |  |
| m (l) | 600 (17.65kb) |  | 722 (17.6 kb) |  | 205 (5.7kb) |  | 803 (5.7 kb) |  | 307 (155kb) |  | 701 (155kb) |  |
| S | 234 |  | 234 |  | 49 |  | 49 |  | 1032 |  | 1032 |  |
| Fc | 0.450 | 0.793 | 0.448 | 0.105 | 0.927 | 0.999 | 0.0588 | 0.001 | 0.630 | 0.972 | 0.632 | 0.17 |
| Gc0 | 23.5 | 0.196 | 46.3 | 0.373 | 73.1 | 0.993 | 2.52 | 0.011 | 14.1 | 0.539 | 28.4 | 0.496 |
| Gc0* | 39.1 | 0.25 | 62.7 | 0.352 | 100.1 | 0.952 | 3.38 | 0.002 | 19.5 | 0.404 | 65.3 | 0.719 |
| Lc0 | 0.0855 | 0.211 | 0.183 | 0.123 | 0.0713 | 0.901 | 0.00558 | 0.001 | 0.0283 | 0.452 | 0.0896 | 0.127 |
| imax | 233 | 0.095 | 555 | 0.911 | 148 | 0.91 | 9 | 0.003 | 197 | 0.726 | 699 | 0.999 |
| gamma10 | 0.260 | 0.109 | 0.352 | 0.075 | 0.545 | 0.973 | 0 | 0.003 | 0.204 | 0.562 | 0.144 | 0.021 |
| gamma10* | 0.439 | 0.138 | 0.481 | 0.037 | 0.750 | 0.873 | 0 | 0.003 | 0.289 | 0.309 | 0.337 | 0.047 |
| C(s.e.) | 3.78 (0.932) |  | 6.73 (1.85) |  | 3.92 (1.63) |  | 0.0785 |  | 12.6 (1.79) |  | 17.5 (3.71) |  |
| ul (x10<sup>-6</sup>/year) | 8.825 |  | 8.825 |  | 2.85 |  | 2.85 |  | 77.5 |  | 77.5 |  |
| tD(my) | 0.428329 |  | 0.762606 |  | 1.375439 |  | 0.02754386 |  | 0.162581 |  | 0.225806 |  |
| (ky) | 428 |  | 763 |  | 1375 |  | 27.5 |  | 163 |  | 225 |  |

To further clarify the population dynamics of *GHRd3* and the non-deleted alleles in YRI and CHB, we applied an approximate Bayesian computation framework developed in a previous study (*15*) to infer the mode and tempo of natural selection. We first simulated data from three different models, the selection on a new mutation, selection on a standing variant, and neutrality, by using four parameters: allele frequency at present (*f*_0_), the onset of natural selection (*t*), selection coefficient (*s*), and allele frequency at *t* (*f_t_*). The details of our simulation conditions are shown in the table below. We then calculated the goodness-of-fit of each model to the observed summaries (**Fig. S6**) as an approximate marginal likelihood (aML) using a kernel density estimate (*61*). Once we identified the best fitting model in terms of approximate Bayes factors (aBFs), we estimated the parameters under the model by kernel ABC (*62*).

| Population | Demography | Prior conditions |  |  |
| --- | --- | --- | --- | --- |
| | | $s$ | $t$ | $f_t$ |
| YRI | Constant ($N = 12,000$) | $\log_{10}U(-3, -0.5)$ | $Log-normal(1600, 1600^2)$ | $U(0.0, 0.2)$ |
| CHB | Population bottleneck and expansion<br>~1600 generations ago: $N = 12,000$<br>1100~1600: $N = 1,200$<br>500~1100: $N = 12,000$<br>500~Present: $N = 120,000$ | | | |

In the YRI population, the standing variation model showed the highest aML. However, this aML was almost comparable to the one under the neutral model (see the table below). The aML calculated for the CHB population was significantly higher in the standing variation model than in the other models, suggesting that the non-deleted allele became advantageous during the expansion into East Eurasia.

To test this hypothesis, we further estimated the onset of natural selection under the standing variation model (see **Fig. 1E**). The estimates showed that selection on the non-deleted allele occurred around 30,000 years ago when this allele was present at 11%. The frequency increased over time with a selection coefficient of 1.13%.

| Population | Models |
| --- | --- |

|  |  | Neutral | Selection on new mutation | Selection on standing variant |
| --- | --- | --- | --- | --- |
| YRI | Log(aML<sup>1</sup>) | -214.687 | -249.968 | -212.995 |
|  | Log<sub>10</sub>(aBF<sup>2</sup>) between a given model and the model with the highest aML | 0.735 | 16.057 | - |
| CHB | Log(aML) | -206.437 | -224.875 | -194.349 |
|  | Log<sub>10</sub>(aBF) between a given model and the model with the highest aML | 5.250 | 13.257 | - |

<sup>1</sup>Approximated marginal likelihood under a given model.
<sup>2</sup>Approximated Bayes factor shows how much one model is better fit to the observed data than the other.

#### Phenotype-wide association studies

One of the single nucleotide variants (rs4073476) upstream of the exon 3 deletion almost perfectly tags *GHRd3* in European populations (R^2^ >0.95). This variant was entered into the GeneATLAS PheWAS database (http://geneatlas.roslin.ed.ac.uk/phewas/) for an association analysis (*16*). This database identifies associations between SNPs and 742 phenotypic traits using the UK Biobank cohort (https://www.ukbiobank.ac.uk/). In parallel, we examined the GWAS database (available through https://atlas.ctglab.nl/PheWAS) among 152 phenotypes., we found that the derived *GHRd3* haplotype is strongly associated with bone mineral density. The next most significant association was height with a nominal p-value < 10^-5^ (**Table S1**). To further the association analysis, another database was utilized (http://www.nealelab.is/uk-biobank/). This database, created by the Neale lab, provides GWAS summary statistics for 4,203 phenotypes from the UK Biobank cohort. Summary statistics were downloaded for males, females, and both datasets combined for the top ten associations identified through the previous GeneATLAS analysis. These summary statistics allowed for sex-specific association analyses.

#### Analysis of *GHRd3* in a cohort of severe acute malnutrition

Using available genotyping platforms from previous work (ref), we analyzed Malawi children with edematous severe acute childhood malnutrition (edematous SAM or ESAM) and non-edematous SAM (NESAM) (*21*). The former is a more severe condition. Specifically, we focused on 2 SNPs flanking the GHR exon 3 deletion, rs4590183, and rs6873545, which are in strong (R^2^>0.9) LD with the *GHRd3* in 1000 Genomes data. However, we found that the LD between these two SNPs was broken (R^2^=∼0.55) in this east-central African cohort. Thus, instead of using these SNPs as a proxy for the deletion, we directly genotyped this cohort for the deletion using PCR primers and conditions described earlier (*6*). Briefly, we used the following primers to genotype both the deleted and non-deleted haplotypes.

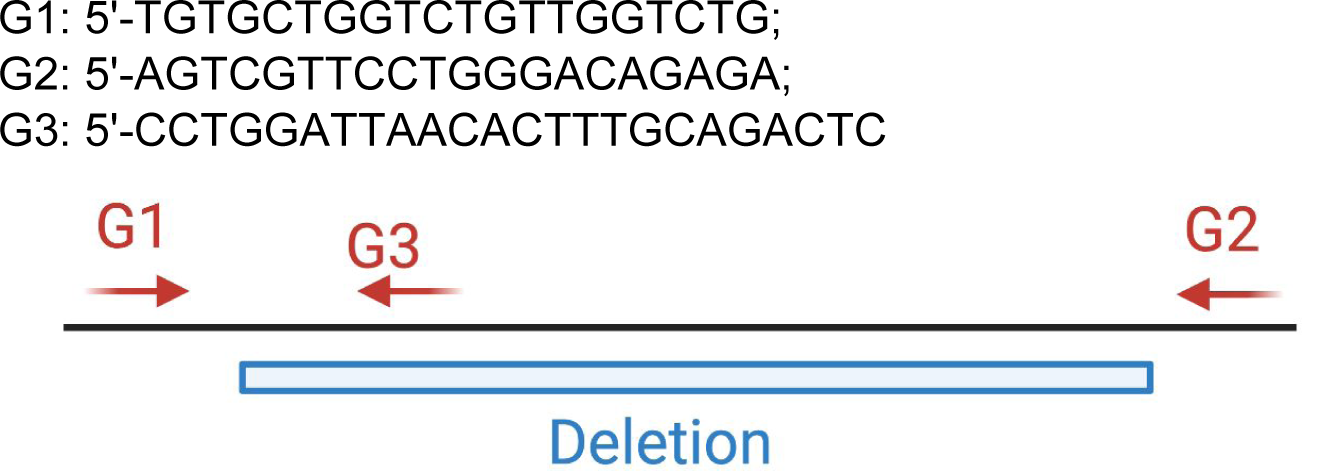

G1: 5’-TGTGCTGGTCTGTTGGTCTG; G2: 5’-AGTCGTTCCTGGGACAGAGA; G3: 5’-CCTGGATTAACACTTTGCAGACTC G1-G3 would not amplify deleted haplotypes but will give a band for non-deleted haplotypes. G1-G2 would not amplify non-deleted haplotypes (too large to amplify) but will give a band for the deleted haplotypes. Overall, we were able to successfully genotype the homozygous and heterozygous deletion with high confidence in 176 samples, consisting of 84 ESAM children (48 males, 36 females) and 92 NESAM children (52 males, 40 females) using these two PCR reactions. We then tested our expectation that GHRd3 is enriched in the less severe form of malnutrition (NESAM). We found strong evidence of an association between the deletion and the ESAM phenotype generally (all individuals, freq in cases vs freq in controls; p<0.01; test), driven by evidence of association in the subset of males (p<0.05).

| Genotype | Phenotype | Gender | Number | Frequency |
| --- | --- | --- | --- | --- |
| Non-Deleted | ESAM | F | 7 | 0.1944 |
| Heterozygous | ESAM | F | 25 | 0.6944 |
| Homozygous Ghrd3 | ESAM | F | 4 | 0.1111 |
| Non-Deleted | NESAM | F | 6 | 0.15 |
| Heterozygous | NESAM | F | 26 | 0.65 |
| Homozygous Ghrd3 | NESAM | F | 8 | 0.2 |
| Non-Deleted | **ESAM** | M | 15 | 0.3125 |
| Heterozygous | **ESAM** | M | 27 | 0.5625 |
| Homozygous Ghrd3 | **ESAM** | M | 6 | 0.125 |
| Non-Deleted | **NESAM** | M | 12 | 0.2308 |
| Heterozygous | **NESAM** | M | 30 | 0.5769 |
| Homozygous Ghrd3 | **NESAM** | M | 10 | 0.1923 |

#### Genome-edited mouse

To model *GHRd3*, we designed sgRNAs that flank both sides (5’ and 3’) of the mouse *Ghr* exon orthologous to the human *GHR* exon 3 with the help of the Gene Targeting and Transgenic Resource at Roswell Park Comprehensive Cancer Center. Specifically, 18 5’ and 11 3’ primers were designed. These primers were screened for common single nucleotide polymorphisms in dbSNP (https://www.ncbi.nlm.nih.gov/snp/) and tested for off target cuts using a standard T7 endonuclease 1 (T7E1) mismatch detection assay (*63*). Based on the results of this assay, the following 4 sgRNA sequences were used for the downstream analysis: mGHR5’.g5 - AATACAATTGGCTAATACCGNGG mGHR5’.g6 - TTGGCTAATACCGAGGTGAGNGG mGHR3’.g10 - TAAGATTTTTAGTGATGTAANGG mGHR3’.g11 - AAACATGACCATTGAATTAANGG These guides were further validated using Bioanalyzer and Qubit for adequate concentrations (∼300ng/uL). C57BL/6N fertilized embryos were harvested at day 0.5 dpc. Fertilized embryos were then injected with guide RNAs and Cas9 into the pronucleus to perform genome editing.

##### In vitro fertilization (IVF) and calorie restriction

To quickly generate a cohort of mice for calorie restriction, d3/d3 males were sent to The Jackson Laboratory for sperm cryopreservation followed by in vitro fertilization of C57BL/6NJ ooyctes and embryo transfer. This resulted in 62 d3 heterozygous mice from which a cohort of 40 mice were generated for the calorie restriction study outlined in **Fig. 2**. The cohorts were kept with their mothers which were fed standard (*Ad libitum*) diets until 1 weaning (∼1month). After weaning, half of the mice were fed 40% calorie restricted chow for another month while the other half were fed *Ad Libitum* diets. All mice were sacrificed at ∼2 months (**Table S2**).

#### Genotyping

The Vindija and Chagyrskaya Neanderthal BAM files were analyzed using the Integrative Genomics Viewer software (v2.3.75) (*64*) to determine if these genomes contained *GHRd3* (https://www.eva.mpg.de/genetics/genome-projects/chagyrskaya-neandertal/home.html?Fsize<u>= 0Svea-Developmental).</u> These results were then added to previous findings regarding the Altai Neanderthal and Denisovan genomes (*9*) **(Fig. S1A)**.

The mice were screened for *GHRd3* using the Kapa Biosystems KAPA2G Fast HS Genotyping kit alongside two sets of primers (**Fig. S8**), following the producer’s protocol. One set of primers leads to amplification if the deletion is present; the other set of primers leads to amplification if the non-deleted, ancestral haplotype is present. For heterozygous individuals, amplification occurs with both sets of primers. For each experiment, water was used as a negative control and a DNA sample from a known heterozygote was used as a positive control.

Deletion Primers: Forward - SM444CEL F (AGAGTACCCAGTGTATGGCCT). Reverse - SM445CEL R (TGCTGTCTGGCACACATGAT) Non-Deletion Primers: Forward SM444CEL F (AGAGTACCCAGTGTATGGCCT). Reverse - SM444CEL R (AGTTCTGTGAGCTGGTGTAGC) In parallel, to validate the successful genome-editing of the mice at the transcriptome level, we conducted a read-depth analysis on BAM files made from the transcriptomes. To create BAM files, FASTQ files were aligned to the GRCm38 (mm10) mouse reference genome using TopHat2 (*65*) and Bowtie2 (*66*). We then observed the read depths of transcripts from exon 3 and exon 4 (the latter for comparison) of the mouse *Ghr* by using the following Samtools command (*67*): samtools depth -q 0 -Q 0 -b mouseGHRex3.bed M3.bam | awk ’{sum+=$3;cnt++}END{print sum/cnt” “sum” “cnt}’> M3.depth Based on the read-depth information, the presence or absence of exon 3 expression was readily detectable (**Fig. S9**).

##### Checking for in vitro fertilization effects

To double-check the potential off-target effects from CRISPR-cas9 gene editing, we sequenced 3 male homozygous GHRd3 mice (0063, 0008, and 0077) to at least 50X coverage on an Illumina Novaseq S1 flowcell. We used BWA (*68*) to align these reads against the GRCm38 reference genome and called single nucleotide and insertion-deletion variants by GATK recommended pipeline (https://gatk.broadinstitute.org/hc/en-us), using the dbSNP dataset as known variable sites. Assuming that the off-target effects that are not proximate to the GHR locus will be eliminated by the multiple backcrossings, we specifically checked 250kb upstream and downstream around the GHR gene (∼1Mb region in total including the GHR gene). In this region, we found 12 variants (2 SNPs and 10 indels) that pass the routine QC thresholds (removing sites with QD < 2.0, QUAL < 30.0, SOR > 4.0, FS > 60.0 or ReadPosRankSum < -8.0) and observed in all 6 haplotypes (i.e., homozygous in all three GHRd3 mice that we sequenced) (**Table S6**). The 2 SNPs are previously reported in dbSNP, thus they are most likely drift effects. Of the 10 indels, 8 of them are expansion or contraction of simple repeat arrays and 2 of them fall in recent retrotransposons (**Table S6**). Thus, it is more likely that these indels are recent mutations that drifted to fixation in our colony, alignment artifacts in our calling pipeline, or false-negatives in existing databases rather than off-target effects of the CRISPR-Cas9 intervention. None of these variants hit any known functional sequences. Overall, there is no reason to think that any off-target effects bias the reported results.

#### Transcriptomics data and analysis

28 to 34-day-old mice were sacrificed (**Table S2**) and liver samples were taken and directly put into RNAlater (Thermo Fisher). They were then sent for RNA extraction and sequencing by GENEWIZ. RNA sequencing was performed via Illumina HiSeq, 2x150bp configuration. Quality control of the obtained sequences was performed using FastQC (Andrews S. FASTQC. A quality control tool for high throughput sequence data. http://www.bioinformatics.babraham.ac.uk/projects/fastqc/. Accessed 7/2019), and the results of all the samples were reviewed by MultiQC (*69*). Adaptor sequences and low quality bases were discarded by Trimmomatic (*70*). Filtered reads were mapped to mouse transcriptome reference (GRCm38) from Ensembl (*71*) and quantified using Kallisto (*72*). The transcripts were merged into genes with tximport and Ensembl BioMart (*73*) Differential expression analyses were performed by DESeq2 (*74*), which calculates the fold change of each gene using the Wald test and a correction for multiple hypotheses. The gene expressions of the samples are also provided in **Table S3**. We then defined genes that were upregulated or downregulated using the adjusted (i.e., multiple hypothesis corrected) *p-*value threshold of 0.05 (Wald test in DeSeq2).

The genes that were upregulated or downregulated in *Ghrd3* mice (**Table S3**) were used against a whole genome background to investigate gene ontology enrichment for all available datasets using ShinyGo (*75*).

#### Lipidomics Analysis

An internal standard mix was prepared in CHCl_3_ containing exogenous lipids *d*_9_ oleic acid (2 μM), *d*_11_ arachidonic acid (2 μM), *d*70 distearoylphosphatidylcholine (2 μM), *d*_31_ C16 sphingomyelin (2 μM), C17 glucosylceramide (2 μM), C57 triacylglycerol (1 μM), C28 diacylglycerol (0.5 μM), dihydrolanosterol (10 μM), and C17 ceramide (0.5 μM).

Lipid extraction and analysis were carried out as described in previous studies (*76, 77*). 28 to 34-day-old mice were sacrificed and serum samples were syphoned. The serum was collected using standard centrifugation methods (*78*) and later stored at -20 degrees. For the LC-MS analysis, serum samples were thawed on ice, gently mixed by pipetting, then 30 μL samples (or water for blank extractions) were transferred to 1 mL cold PBS contained in a 2-dram glass vial on ice. Samples were then vortexed three times for 3 seconds to mix and a 30 μL aliquot was taken for protein measurement using a Coomassie assay. The relative standard deviation in protein concentration among the samples was 5%. 1 mL of methanol, 1.8 mL of CHCl_3_, and 200 μL of internal standard mix were added to the remainder. All solutions were kept cold on ice. Samples were vortexed for 10 seconds, followed by 1 min on ice, three times. Samples were then centrifuged for 30 minutes at 3000 rcf at 4°C. The bottom (CHCl_3_) layer was transferred to a 1 dram glass vial and kept on ice. The upper layer was re-extracted with 2 mL of additional CHCl_3_ and the two CHCl_3_ layers were combined. 3.75 mL of the combined CHCl_3_ layers were transferred to a new 1 gram vial and the solvent was removed by rotary evaporation.

Samples were resuspended in 150 μL of CHCl_3_ spiked with ^13^C_18_ oleic acid (2 μM), C39 triacylglycerol (1 μM), and C6 ceramide (0.5 μM). Samples were analyzed by LC-MS using an Agilent Infinity 1260 HPLC/ Agilent 6530 Jet Stream ESI-QToF-MS system. Mobile phase A was composed with 95% H_2_O and 5% methanol. Mobile phase B was composed with 60% isopropanol, 35% methanol, and 5% water. Samples were analyzed in positive ionization mode using 0.1% (v/v) formic acid and 5 mM ammonium formate as additives in the mobile phases with separation performed using a Luna C5 column (5 µm C5 100 Å, 50 x 4.6 mm) with a C5 guard cartridge. Samples were analyzed in negative ionization mode using and 0.1% ammonium hydroxide as an additive in the mobile phases with separation performed using a Gemini C18 column (5 µm C18 110 Å, 50 x 4.6 mm) with a C18 guard cartridge. 5 μL of the resuspended sample was injected for analysis. The LC method began with 5 min of 0% B at 0.1 mL/min, then the flow rate was increased to 0.5 mL/min and mobile phase gradient began with 0% B to 100% B over 60 min. 100% B was maintained for 7 min, then switched to 0% B for 8 min. LC-MS data analysis was performed using the Agilent MassHunter Qualitative Analysis software (v. B.06.00). LC-MS grade methanol and isopropanol were obtained from Millipore Sigma; LC-MS grade chloroform was obtained from Honeywell. Internal standards for LC-MS were obtained from Avanti Polar lipids, with the exceptions of ^13^C_18_ oleic acid (Cambridge Isotope laboratories) and C39, C57 TAGs (Millipore Sigma). LC columns were obtained from Phenomenex. Coomassie protein assay kits were obtained from Thermo Scientific.

The lipid levels of the samples are provided in **Table S5**.

### Supplementary tables

Table S1. SNPs in Linkage disequlibrium with GHRd3 and PheWAS results

Table S2. Sample information and weight data

Table S3. Transcriptome data

Table S4. GO analysis results

Table S5. Lipidomics results

